# The *Caenorhabditis elegans* bacterial microbiome influences microsporidia infection through nutrient limitation and inhibiting parasite invasion

**DOI:** 10.1101/2024.06.05.597580

**Authors:** Hala Tamim El Jarkass, Stefanie Castelblanco, Manpreet Kaur, Yin Chen Wan, Abigail E. Ellis, Ryan D. Sheldon, Evan C. Lien, Nick O. Burton, Gerard D. Wright, Aaron W. Reinke

**Affiliations:** Department of Molecular Genetics, University of Toronto, Toronto, ON, Canada; Michael G. DeGroote Institute for Infectious Disease Research & David Braley Centre for Antibiotic Discovery. McMaster University, Hamilton, ON, Canada; Mass Spectrometry Core, Van Andel Research Institute. Grand Rapids, MI USA; Department of Metabolism and Nutritional Programming, Van Andel Research Institute. Grand Rapids, MI USA

## Abstract

Microsporidia are eukaryotic obligate intracellular parasites that infect most animals including humans. To understand how the microbiome can impact microsporidia infection, we tested how bacterial isolates that naturally occur with *Caenorhabditis elegans* influence infection by the microsporidian *Nematocida parisii*. Nematodes exposed to two of these bacteria, *Chryseobacterium scopthalmum* and *Sphingobacterium multivorum*, exhibit reduced pathogen loads. Using untargeted metabolomics, we show that unsaturated fatty acid levels are disrupted by growth on these bacteria and that supplementation with the polyunsaturated fatty acid linoleic acid can restore full parasite growth in animals cultured on *S. multivorum*. We also found that two isolates, *Pseudomonas lurida* and *Pseudomonas mendocina,* secrete molecules that inactivate *N. parisii* spores. We determined that *P. lurida* inhibits *N. parisii* through the production of massetolides. We then measured 53 additional *Pseudomonas* strains, 64% of which significantly reduced *N. parisii* infection. A mixture of *Pseudomonas* species can greatly limit the amount of infection in *C. elegans* populations over many generations. Our findings suggest that interactions between bacteria and *N. parisii* are common and that these bacteria both modulate host metabolism and produce compounds that inhibit microsporidia infection.

## Introduction

Organisms are tightly connected with the microbial entities that exist both within and around them. Some members of the microbiome act as pathogens, causing disease, whereas others play important roles in host health (Hou et al. 2022). Beneficial members of the microbiome can protect against pathogens through limiting host resources, secretion of antimicrobial compounds, or stimulating host immunity (Spragge et al. 2023; Liu et al. 2020; Chevrette et al. 2019). These beneficial microbes can both protect the host and influence how pathogens evolve (Hoang, Read, and King 2024; Ford et al. 2016). Understanding these mechanisms of host protection can lead to harnessing bacteria to prevent infectious disease (Jin Song et al. 2019).

The model organism *Caenorhabditis elegans* has become a powerful system to mechanistically study complex interactions between a host, beneficial microbes, and pathogens (Schulenburg and Félix 2017). This invertebrate is commonly isolated from rotting plant matter containing over 250 distinct bacterial isolates consisting of mostly Gammaproteobacteria and Bacteroidetes (Frézal and Félix 2015; F. Zhang et al. 2017). These associated bacteria predominantly act as a food source for *C. elegans*, though 20% of these bacterial isolates can be pathogenic (Samuel et al. 2016). A community effort combining sampling efforts from distinct geographical locations resulted in the generation of a common set of bacteria that frequently co-occur with this nematode in the wild. This resulting bacterial collection, known as CeMbio, is composed of 12 different species that represent ∼60% of the known diversity of *C. elegans*-associated bacteria (Dirksen et al. 2020). Members of this collection can defend against bacterial and viral pathogens such as *Pseudomonas mendocina* MSPm1 protecting against *Pseudomonas aeruginosa* through activation of the p38 Mitogen-activated protein kinase pathway, *Pseudomonas lurida* MYb11 producing the antimicrobial compound massetolide E to inhibit *Bacillus thuringiensis* infection, and JUb44 *Chryseobacterium scopthalmum* providing resistance to Orsay virus infection that is independent of known antiviral pathways (Kissoyan et al. 2019; Gonzalez and Irazoqui 2023; Montalvo-Katz et al. 2013; Vassallo et al. 2023; González and Félix 2024).

Microsporidia are ubiquitous obligate intracellular fungal parasites that infect most types of animals (Murareanu et al. 2021; Han and Weiss 2017; Wadi and Reinke 2020). These parasites can cause disease in humans as well as agriculturally important animals such as fish, shrimp, honey bees, and silkworms (Stentiford et al. 2016; Han and Weiss 2018; Bojko et al. 2022). The presence of microsporidia in the microbiome has been extensively studied, with reported prevalence rates of over 60% in animal populations (Ruan et al. 2021; Trzebny et al. 2020; Shen et al. 2019; Martín-Hernández et al. 2018). Possessing the smallest known eukaryotic genome sizes, these parasites have lost many metabolic enzymes and are extremely host dependent, utilizing a variety of transporters to uptake nutrients and ATP (Nakjang et al. 2013; Dean et al. 2018; Heinz et al. 2014; Tsaousis et al. 2008; Major et al. 2019; Wadi et al. 2023). The host microbial diet can impact microsporidia such as bacterially-produced vitamin B12 providing tolerance to infection (Willis et al. 2023). Bacteria can also produce molecules that inhibit microsporidia and bacteria engineered to produce double stranded RNA protect against microsporidia infection (Q. Huang et al. 2023; Lang et al. 2023; Porrini et al. 2010; Tersigni et al. 2024).

Microsporidia are common parasites of *C. elegans* in nature, and the most frequently observed species is *Nematocida parisii* (Troemel et al. 2008; G. Zhang et al. 2016; Luallen et al. 2016; Reinke et al. 2017; Wadi et al. 2023). Infection begins when *C. elegans* ingest dormant *N. parisii* spores. These spores contain a unique invasion apparatus called the polar tube. When spores come into contact with a host cell, the polar tube rapidly emerges and is thought to inject the intracellular components of the spore, the sporoplasm, inside of the target host cell, which can occur as early as two minutes after being exposed to spores (Tamim El Jarkass and Reinke 2020; Luallen et al. 2016). In approximately 12-18 hours, meronts will begin to form, resulting in the formation of mature dormant spores as early as 48 hours post-infection (Balla et al. 2016). *N. parisii* infections in the wild happen within the context of the large collection of bacterial species that are associated with *C. elegans*, but how the bacterial diet influences *N. parisii* growth or if these bacteria produce molecules that inhibit microsporidia infection is unknown.

To understand how the *C. elegans* microbiome impacts *N. parisii* infection, we utilized the CeMbio collection of bacteria. We show that *Sphingobacterium multivorum* BIGb0170 and *Chryseobacterium scopthalmum* JUb44 delay *N. parisii* growth through nutrient limitation. We show that growth on *S. multivorum* BIGb0170 causes disruption of unsaturated fatty acids within *C. elegans,* and that supplementation of linoleic acid can restore *N. parisii* growth. We show that linoleic acid is a limiting factor for *N. parisii* to complete its lifecycle as an infected *C. elegans* mutant that is unable to make this polyunsaturated fatty acid experiences delayed differentiation into spores. We then show that *P. lurida* MYb11 and *P. mendocina* MSPm1 secrete distinct compounds that inhibit *N. parisii* spores. We tested an additional 53 *C. elegans*-associated *Pseudomonas* isolates, demonstrating that 64% of them can inhibit *N. parisii*. We also show that a mixture of eight *Pseudomonas* species reduced *N. parisii* infection by 80% and that this protection lasts at least 10 generations. Altogether, our study shows how the *C. elegans* microbiome can impact *N. parisii* infection by disrupting unsaturated fatty acid availability and that many *Pseudomonas* species secrete molecules that inactivate *N. parisii* spores.

## Results

### Members of the *C. elegans* microbiome reduce *C. elegans* feeding and *N. parisii* growth through nutritional deficiency

To investigate how growth on different bacterial isolates impacts *N. parisii* infection, we exposed wild-type N2 L1 stage *C. elegans* to individual lawns of CeMbio bacterial strains for 72 hours. All CeMbio strains were cultured at 25°C and as a control we included the typical lab diet of *Escherichia coli* OP50 grown at the routine temperature of 37°C or at 25°C (see methods). These young adult animals were then exposed to 48-hours of continuous infection with *N. parisii* spores, fixed, and stained with direct yellow 96 (DY96), a dye that binds to the chitin-containing spores **(Figure 1a)**. *N. parisii* growth was hindered when nematodes were grown on 7 of the 11 CeMbio strains as determined by a reduction in the fraction of worms producing spores **(Figure 1b)**.

**Figure 1.**
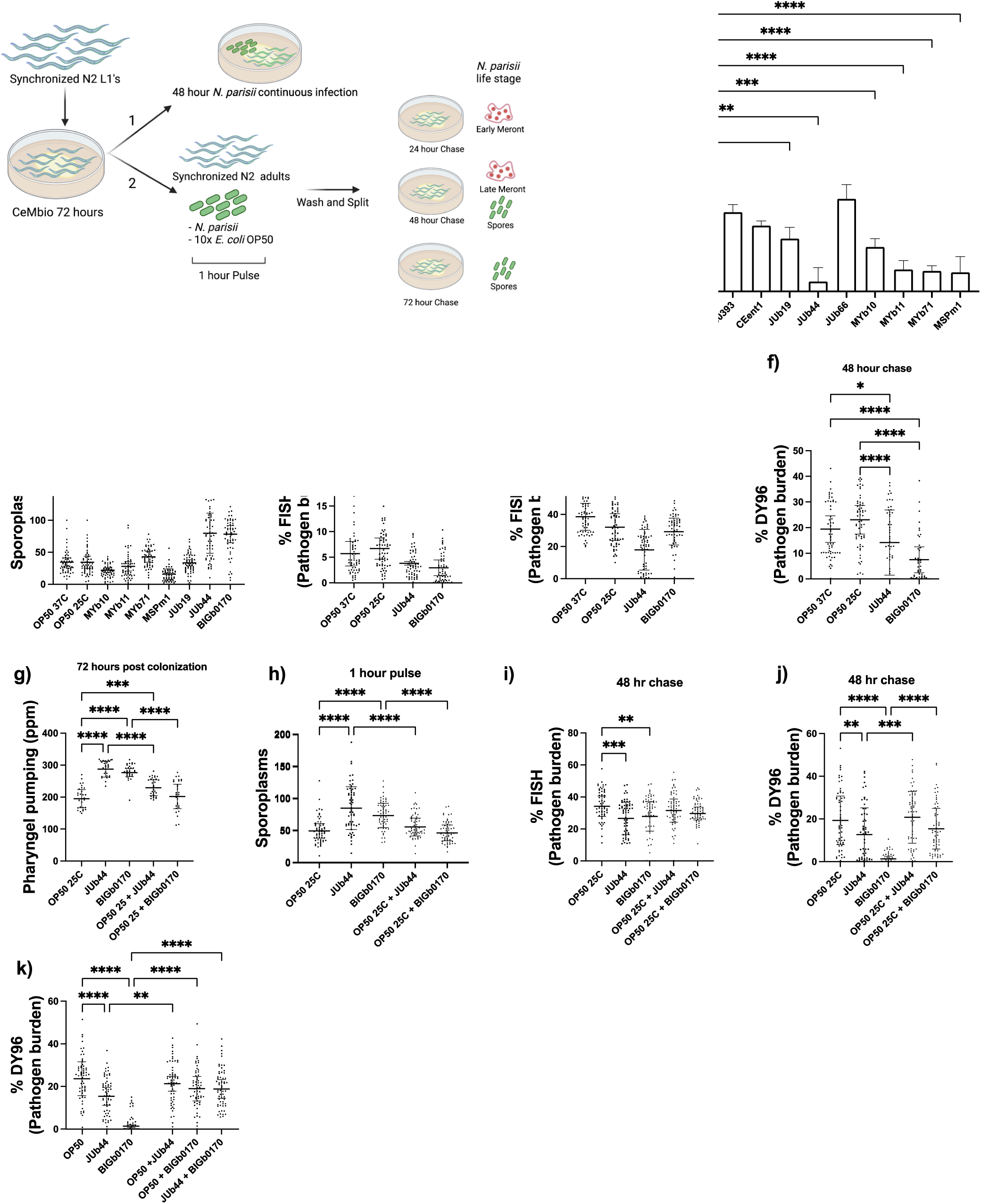
Growth on *C. scopthalmum* JUb44 and *S. multivorum* BIGb0170 results in decreased *N. parisii* infection burden. (a) Schematic depicting the workflow involved in the colonization of nematodes with CeMbio strains and the experimental pipeline. Synchronized L1 animals were grown on individual bacterial strains for 72 hours. Synchronized adult animals were (1) continuously infected with *N. parisii* on CeMbio lawns or (2) exposed to a 1-hour pulse, followed by a 24 or 48-hour chase. Sporoplasms and meronts can be detected at earlier time points using FISH and spores can be detected at later time points using DY96. (b) The fraction of worms displaying spores 48 hpi. (c-f) 72-hour old adults were pulse infected with *N. parisii* and *E. coli* OP50 for one hour and fixed at 1 hpi (c) 24 hpi (d) or 48hpi (e,f) and stained with FISH probes and DY96 and amount of pathogen load was quantified. (g-j) Bacterial strains were used in isolation or in a 1:1 ratio with *E. coli* OP50 and nematodes were grown for 72 hours. (g) Quantification of pharyngeal pumps per minute. 72-hour colonized nematodes were pulse infected for one hour and sporoplasms (h), meronts (i) and spores (j) quantified. (k) 72-hour old adults grown on the indicated bacteria were pulse infected for one hour and spores quantified. Data is from three independent replicates of 100 (b), 15-20 (c-f,h-k) or 10 worms each (g). Mean ± SD represented by horizontal bars. P-values determined via one-way ANOVA with post hoc. Significance defined as * p < 0.05, ** p < 0.01, *** p < 0.001, **** p < 0.0001.

The first step of microsporidia infection is the invasion of host cells by microsporidian sporoplasms. To determine if invasion is impacted by growth on CeMbio strains, we exposed *C. elegans* adults grown on bacterial strains for 72-hours to *N. parisii* spores. After 1 hour of exposure, animals were fixed and stained with an *N. parisii*-specific 18S rRNA Fluorescence *in situ* Hybridization (FISH) probe. Nematodes grown on *P. mendocina* MSPm1 had a reduction in the number of sporoplasms whereas animals grown on *S. multivorum* BIGb0170 or *C. scopthalmum* JUb44 displayed a significant increase in the number of sporoplasms **(Figure 1c)**. Although animals grown on *S. multivorum* BIGb0170 and *C. scopthalmum* JUb44 were initially more infected, they produced fewer spores later **(Figure 1b).**

To understand how growth on *S. multivorum* BIGb0170 or *C. scopthalmum* JUb44 results in decreased *N. parisii* proliferation, we performed a pulse-chase assay, monitoring how *N. parisii* infection progresses with time **(Figure 1a)**. 72-hour old *C. elegans* adults were exposed to a one-hour pulse infection with *N. parisii* and un-ingested spores were washed away prior to placing the animals back on the corresponding lawns of bacteria. Animals were then washed and fixed at various time points and stained using a FISH probe and DY96. At both 24-hours post infection (hpi) and 48 hpi, a reduction of meronts was observed, and at 48 hpi a reduction in spores was observed **(Figure 1d-f)**. L1 stage *C. elegans* grown on *S. multivorum* BIGb0170 or *C. scopthalmum* JUb44 for only 24 hours prior to infection did not display increased invasion or decreased pathogen proliferation **(Figure S1a-d)**.

*S. multivorum* BIGb0170 and *C. scopthalmum* JUb44 were previously reported to result in developmental delay of *C. elegans*, suggesting they may not be optimal food sources (Dirksen et al. 2020). Poor quality diets can influence nematode feeding behaviors, including pharyngeal pumping (Avery and You 2012; Shtonda and Avery 2006; Scholz et al. 2016). Nematodes grown on *S. multivorum* BIGb0170 or *C. scopthalmum* JUb44 for 72 hours displayed significantly increased levels of pharyngeal pumping **(Figure 1g)**. These findings indicate that nematode feeding behaviour is intensified by these bacterial species. To assess if this behavioural change was a result of altered nutritional quality, we grew nematodes on lawns of either bacterial species mixed with *E. coli* OP50 in a 1:1 ratio, which resulted in significantly reduced pharyngeal pumping rates **(Figure 1g)**. Animals grown on these mixed lawns also resulted in less initial *N. parisii* invasion and an increase in pathogen burden relative to infected nematodes grown on either *S. multivorum* BIGb0170 or *C. scopthalmum* JUb44 only **(Figure 1h-j)**. To determine if these two bacteria could complement each other, we grew nematodes on mixed lawns containing a 1:1 ratio of *C. scopthalmum* JUb44 and *S. multivorum* BIGb0170. The mixture resulted in wild-type levels of *N. parisii* spore production **(Figure 1k)**. Together, our results suggest that more active pharyngeal pumping results in increased *N. parisii* invasion and a nutritional component of the *E. coli* OP50 diet may be missing from *S. multivorum* BIGb0170 and *C. scopthalmum* JUb44.

### Linoleic acid is a limiting factor for *N. parisii* growth

To determine if growth on *S. multivorum* BIGb0170 and *C. scopthalmum* JUb44 altered the metabolites present in *C. elegans*, we performed untargeted metabolomics. We grew animals for 72 hours on either *E. coli* OP50, *S. multivorum* BIGb0170, or *C. scopthalmum* JUb44. This analysis revealed that animals grown on either *S. multivorum* BIGb0170 or *C. scopthalmum* JUb44 had altered levels of three polyunsaturated (PUFA) lipid classes: triglycerides (TG), phosphatidylcholine (PC), and phosphatidylethanolamine (PE) relative to animals grown on *E. coli* OP50 **(Figure 2a, S2a,b)**. Animals grown on *C. scopthalmum* JUb44 had largely reduced TG levels whereas animals grown on *S. multivorum* BIGb0170 contained classes of TGs that were either reduced or elevated compared to animals grown on *E. coli* OP50. Animals grown on *C. scopthalmum* JUb44 had significantly fewer TGs with acyl chains lengths less than 17 whereas those grown on *S. multiovrum* BIGb0170 had significantly more **(Figure 2b)**.

**Figure 2.**
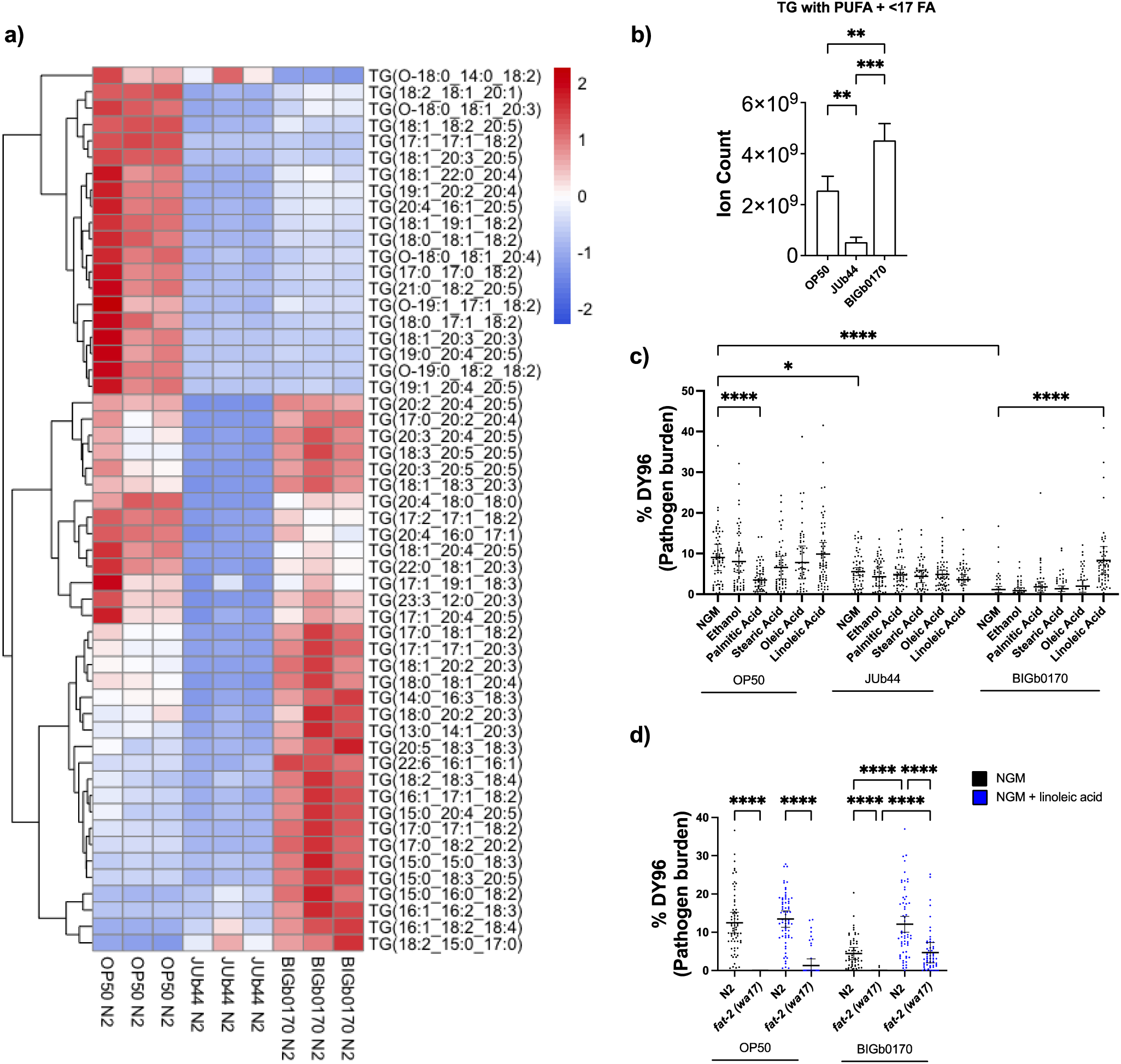
Linoleic acid supplementation rescues *N. parisii* growth on *S. multivorum* BIGb0170. (a) A heat map depicting the abundance of triglycerides (TG) containing a polyunsaturated fatty acid (PUFA) acyl chain in N2 animals grown on *E. coli* OP50, *C. scopthalmum* JUb44 or *S. multivorum* BIGb0170 from three independent replicates. Scale bar represents Z-score values. (b) Ion counts of PUFA triglycerides containing fewer than 17 carbons in N2 animals grown on the different bacterial diets. (c) Various mono- and polyunsaturated fatty acids were supplemented in NGM media when nematodes were colonized with bacterial species denoted on the X-axis and spores quantified. (d) N2 or *fat-2 (wa17)* animals were grown on plates with (blue) or without (black) linoleic acid. The bacterial diet is denoted with a solid line below the X-axis. Data is from three independent replicates of 20 worms each (c-d). Mean ± SD represented by horizontal bars. P-values determined via one-way ANOVA with post hoc. Significance defined as * p < 0.05, ** p < 0.01, *** p < 0.001, **** p < 0.0001.

To test whether fatty acids are necessary for microsporidia growth on *S. multivorum* BIGb0170 and *C. scopthalmum* JUb44, we supplemented bacterial diets with various fatty acids. Supplementation of palmitic (C16:0), stearic (C18:0), oleic (C18:1), or linoleic (C18:2) acid revealed that only the polyunsaturated linoleic acid was able to restore *N. parisii* growth in animals grown on *S. multivorum* BIGb0170. We did not observe an effect of fatty acid supplementation on *N. parisii* growth on animals grown on *C. scopthalmum* JUb44 **(Figure 2c)**. The abundance of C18:2-containing TGs differed between animals grown on JUb44 and BIGb0170 whereas C18:2-containing PE and PCs were generally lower overall **(Figure S2c-e)**.

To test the importance of linoleic acid on *N. parisii* proliferation we used a mutant of *fat-2*, which encodes a Δ12 desaturase and plays an essential role in the conversion of oleic acid (C18:1) to linoleic acid (C18:2). Animals lacking FAT-2 contain depleted amounts of linoleic acid (Watts and Browse 2002). When *fat-2 (wa17*) animals were grown on *E. coli* OP50 or *S. multivorum* BIGb0170 for 72 hours, we observed a complete block in *N. parisii* sporulation 48 hours later **(Figure 2d)**. To confirm that this phenotype was due to a lack of linoleic acid, we performed the same experiment with supplementation of linoleic acid. *fat-2 (wa17)* animals displayed improved *N. parisii* sporulation on both bacterial sources when linoleic acid was supplemented. Linoleic acid supplementation also improved *fat-2 (wa17)* fitness as seen by an increase in the number of embryos after 72 hours of growth on both *E. coli* OP50 and *C. scopthalmum* BIGb0170 **(Figure S3a)**.

To determine if a lack of linoleic acid impacted *N. parisii* growth over time, we performed pulse-chase experiments. *fat-2 (wa17)* animals grown on either *E. coli* OP50 or *C. scopthalmum* BIGb0170 for 48 hours displayed reduced levels of meronts compared to N2 animals **(Figure S3b)**. At 48 hpi, sporulation was also greatly reduced in the *fat-2* mutant **(Figure S3c)**. However, at 72 hpi, *fat-2 (wa17)* animals displayed higher levels of sporulation than observed at 48 hours on either diet, though still lower than in N2 animals **(Figure S3d)**. Together these results suggest that linoleic acid is a limiting nutrient for efficient growth of *N. parisii* in *C. elegans*.

As animals grown on *C. scopthalmum* BIGb0170 resulted in increased sporoplasm invasion, we investigated if linoleic acid was responsible for this phenotype. We measured the number of sporoplasms in N2 and *fat-2 (wa17)* animals and observed no differences, and also observed no differences if the media was supplemented with linoleic acid **(Figure S4a)**. Oleic acid has previously been reported to be important for feeding behavior in *C. elegans* (Hyun et al. 2016). To determine if supplementation with this lipid could rescue the increased invasion, we performed invasion experiments, observing that oleic acid, but not linoleic acid, reduced sporoplasm levels when worms were grown on *C. scopthalmum* BIGb0170 **(Figure S4b)**. Supplementation with oleic acid also reduced the increase in pharyngeal pumping caused by *C. scopthalmum* BIGb0170 **(Figure S4c)**. Supplementation with oleic acid did not have an effect on the increased sporoplasms or pharyngeal pumping when animals were grown on *C. scopthalmum* JUb44 **(Figure S4b,c).** Together this data shows that oleic acid can rescue the feeding behaviour of animals grown on *C. scopthalmum* BIGb0170 and provides further support that the nutrient deficiency is different between these two Bacteroidetes bacteria.

### Growth on *P. lurida* MYb11 improves *C. elegans* fitness and reduces *N. parisii* proliferation

In addition to providing nutrients to hosts, bacteria also can produce compounds that can protect against infection. Like many microsporidia, *N. parisii* infection reduces embryo formation and overall progeny produced (Willis et al. 2021). To determine how *C. elegans* is impacted during *N. parisii* infection while feeding on these diets, we pulse-infected animals for one hour with *N. parisii* and *E. coli* OP50 and placed infected animals onto individual CeMbio lawns **(Figure 3a).** This approach allows us to rule out any effects that the CeMbio diet may have on *N. parisii* invasion. Growth on most of the CeMbio strains in the absence of infection resulted in the ability of almost all animals to become gravid (containing embryos) 72 hours post L1, except for *S. multivorum* BIGb0170, which is consistent with previous findings (Samuel et al. 2016; Dirksen et al. 2020) **(Figure 3b).** However, when exposed to *N. parisii,* nematodes grown on *P. lurida* MYb11 were ∼4-fold more gravid than those grown on *E. coli* OP50, with population gravidity near that of uninfected animals. We also observed that *Comamonas piscis* BIGb0172 displayed a moderate increase in fitness and those grown on *S. multivorum* BIGb0170 and *C. scopthalmum* JUb44 displayed reduced fitness **(Figure 3b)**.

**Figure 3.**
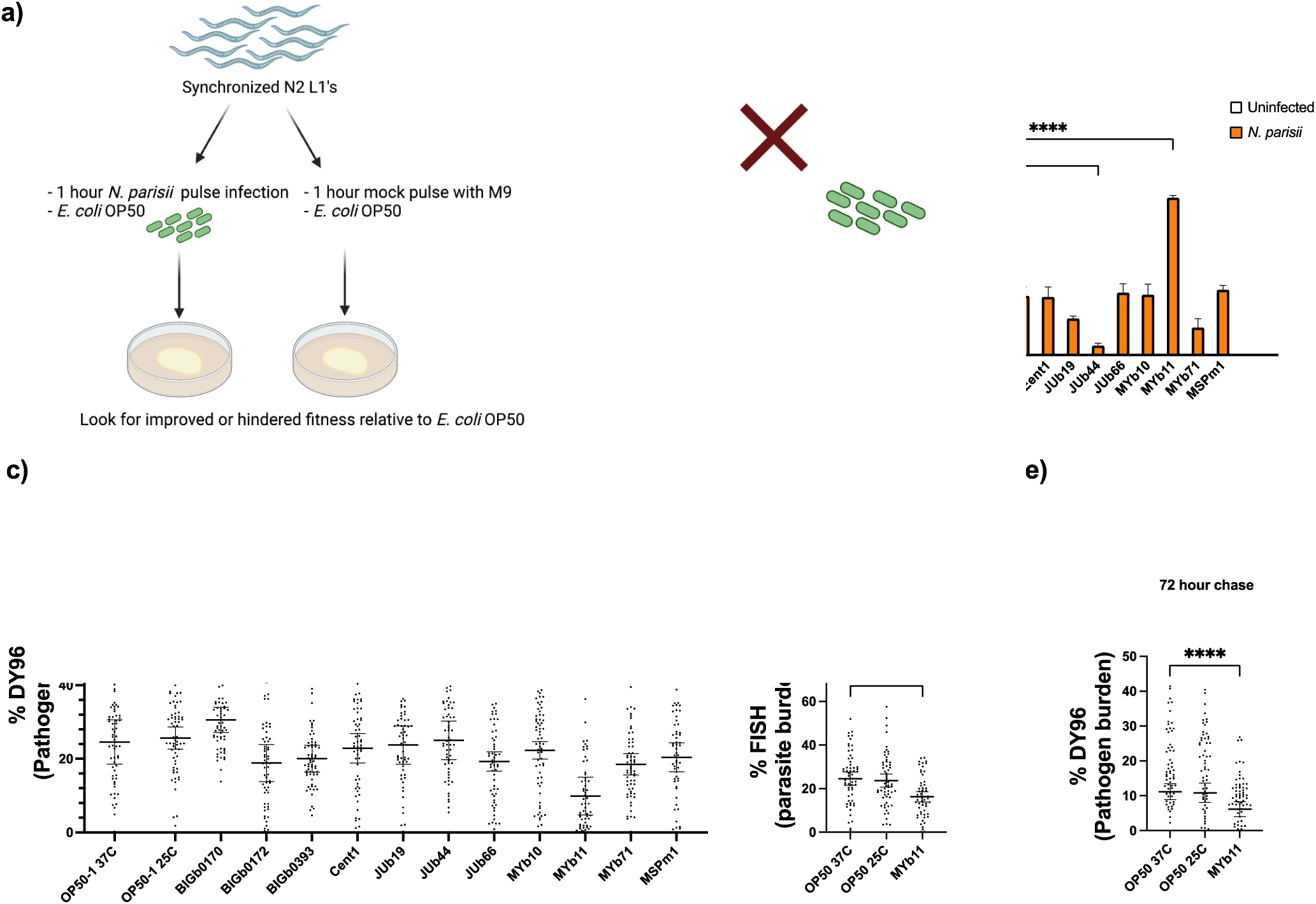
Growth on *P. lurida* MYb11 inhibits *N. parisii* proliferation. (a) Schematic depicting the experimental pipeline. Synchronized N2 L1 animals were either infected with *N. parisii* or mock treated for 1 hour on *E. coli* OP50-1 prior to washing and plating onto various CeMbio strains. 72 hours later, population fitness (b) and pathogen load (c) were quantified. (d-e) Synchronized N2 L1s were pulse infected with *N. parisii* for 1 hour on *E. coli* OP50-1 prior to washing and splitting the populations onto *P. lurida* MYb11 seeded plates. Animals were fixed and then stained with an *N. parisii* 18S RNA fish probe at 48 hpi (d) or DY96 72 hpi (e). Data is from three independent replicates of at least 100 (b) or 15-20 (c-e) worms each. Mean ± SD represented by horizontal bars. P-values determined via one-way ANOVA with post hoc. Significance defined as * p < 0.05, ** p < 0.01, *** p < 0.001, **** p < 0.0001.

We also determined the impact of these diets on *N. parisii* growth. Culturing of nematodes on *P. lurida* MYb11 resulted in ∼2-fold decrease in pathogen load. We also observed that growth on *C. piscis* BIGb0172 and *Ochrobactrum pecoris* MYb71 resulted in a small reduction in pathogen load, and growth on *S. multivorum* BIGB0170 resulted in a small increase in pathogen load **(Figure 3c).** To monitor infection progression on *P. lurida* MYb11, we pulse infected N2 L1 animals for one hour, with 48- (late meront) and 72-hour (spores) chase timepoints. Infected N2 animals grown on *P. lurida* MYb11 consistently displayed reduced pathogen loads throughout development **(Figure 3d,e).** Lastly, to assess if the contributions of the *P. lurida* MYb11 diet were only advantageous at specific timepoints during infection, animals were grown on either *E. coli* OP50 or *P. lurida* MYb11 for varying amounts of time. When animals were exposed to *P. lurida* MYb11 for at least 24 hours from either L1 (0 hours) or L2 (24 hours old), decreased pathogen loads and increases in gravidity were observed. This suggests that the protective effects of the *P. lurida* MYb11 diet are conferred within the first 24 – 48 hours of infection **(Figure S5a,b).**

Previously, it was shown that bacteria producing vitamin B12 induce developmental acceleration in *C. elegans* and several of the CeMbio isolates, including *P. lurida* MYb11, are predicted to contain pathways for vitamin B12 production (MacNeil et al. 2013; Watson et al. 2014; Dirksen et al. 2020). We have previously shown that parents exposed to vitamin B12 produce progeny that have enhanced tolerance to *N. parisii* (Willis et al. 2023). To determine if vitamin B12 is produced in *P. lurida* MYb11, we monitored the expression levels of acyl-CoA dehydrogenase *acdh-1*, a dietary sensory of B12 deficiency (Watson et al. 2014). When *Pacdh-1::gfp* L1 animals were grown on lawns of *Comamonas aquatica* DA1877, a bacterial species known to produce vitamin B12, or *P. lurida* MYb11, GFP expression was repressed **(Figure S6a)**. To determine if the vitamin B12 produced by *P. lurida* MYb11 influenced the enhanced fitness we observed during *N. parisii* infection, we tested mutants in two pathways that utilize vitamin B12: the methionine/SAM cycle *(metr-1)* and the propionyl-CoA breakdown pathway *(mmcm-1)*. Although infected N2 and *mmcm-1(ok1637)* animals display increased fitness when grown on *C. aquatica* DA1877 and *P. lurida* MYb11, only worms grown on *P. lurida* MYb11 display reduced pathogen loads. The *metr-1(ok521)* animals grown on *C. aquatica* DA1877 or *P. lurida* MYb11 no longer display increased fitness, but these mutant animals reared on lawns of *P. lurida* MYb11 still display reduced pathogen loads **(Figure S6b,c).** This supports a model where the enhanced fitness is mediated through production of vitamin B12, but the resistance to infection is not.

### Conditioned media from CeMbio isolates attenuate *N. parisii* spore infectivity

In addition to inhibiting microsporidia proliferation, it is possible that bacteria could make compounds that inhibit microsporidia spores. To investigate if the CeMbio species target *N. parisii,* we incubated spores in conditioned media from individual bacterial species and infected synchronized N2 L1 animals for 72 hours **(Figure 4a)**. *N. parisii* spores incubated in the supernatants of *P. lurida* MYb11, *P. mendocina* MSPm1, and *C. piscis* BIGb0172 were less infective, as indicated by increased proportions of gravid adult animals **(Figure 4b)**. To assess how these supernatants may influence *N. parisii* infectivity, we performed a one-hour pulse infection with supernatant-treated spores to invasion phenotypes **(Figure 4a)**. Microsporidia invasion can be broken down into three main stages: spore ingestion, spore firing, and sporoplasm deposition. A perturbation in any of these three processes would greatly improve host fitness. The number of spores present in the nematode gut were significantly lower under *P. lurida* MYb11 and *P. mendocina* MSPm1 treatment conditions **(Figure 4c)**. Next, we quantified the fraction of fired spores, revealing no significant differences between the different treatment conditions **(Figure 4d)**. Lastly, we quantified the number of intracellular sporoplasms within the nematode intestinal cells. We saw a significant reduction in the number of sporoplasms under *P. lurida* MYb11, *P. mendocina* MSPm1, and *C. piscis* BIGb0172 treatment conditions **(Figure 4e),** with the strongest effect being observed with the two *Pseudomonas* species. The activity of the conditioned media is likely acting directly on the spores, as *N. parisii* spores treated with media and then washed are still less infective, as seen by improved host fitness and reduced pathogen burdens in infected animals under these conditions **(Figure S7a,b).**

**Figure 4.**
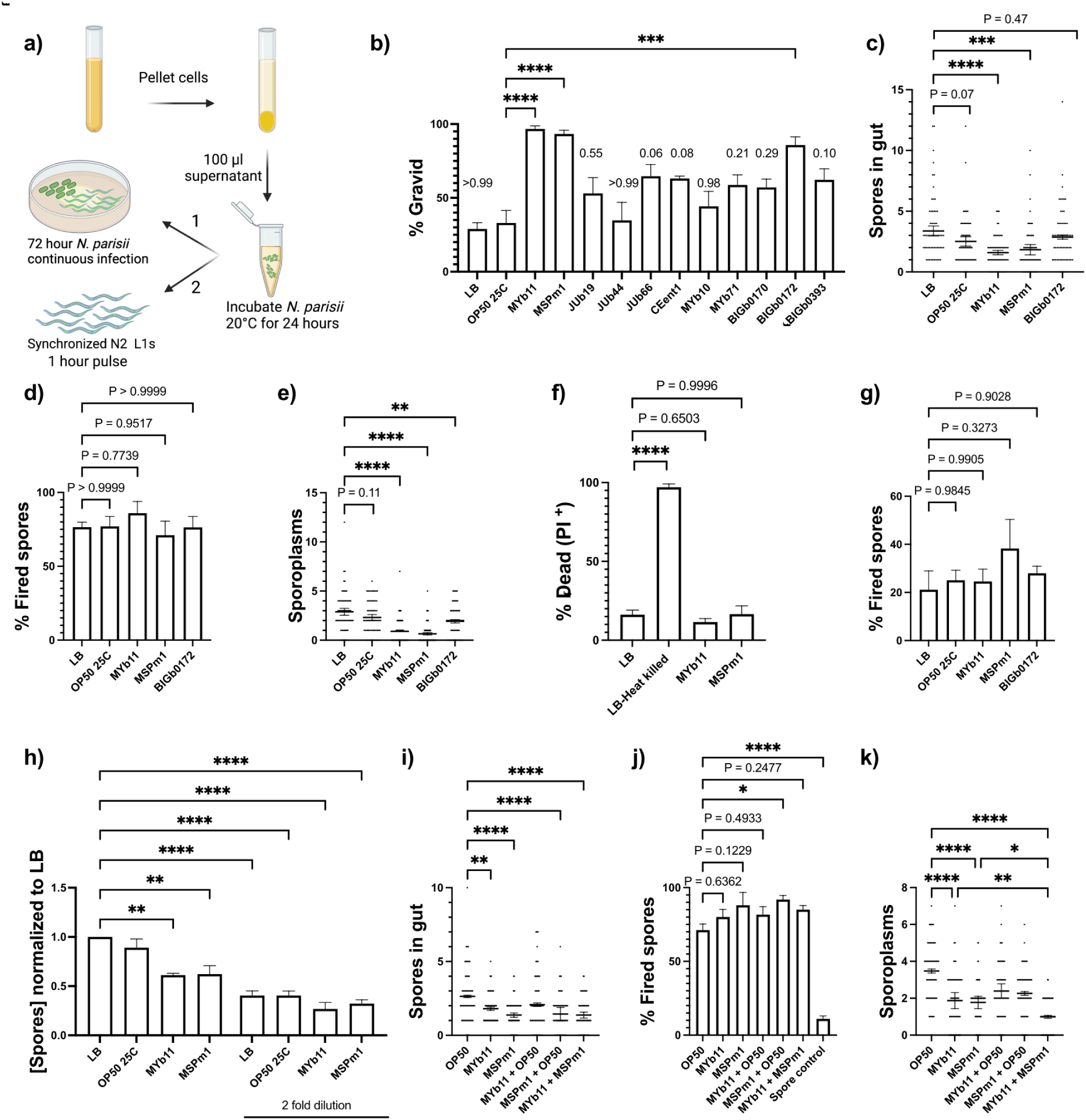
*P. lurida* MYb11 and *P. mendocina* MSPm1 secrete molecules which cause *N. parisii* spore destruction. (a) Supernatants from various CeMbio strains of interest were incubated with *N. parisii* spores for 24 hours at 21°C. Synchronized N2 L1 animals and *E. coli* OP50 were then added to the spore-supernatant mixtures and plated on a 6-cm unseeded NGM plate for 72 hours (1) (b) or 1 hour (2) (c-e, i-k). 72 hpi population fitness was quantified (b). Values above bars represent p-values relative to *E. coli* OP50. (c-e) Pre-incubated spores were used to infect N2 L1’s for one hour as in (a)-2. Pathogen invasion was assessed. The number of spores per animal (c), the percentage of fired spores (d) and the number of sporoplasms (e) are displayed. (f-h) *N. parisii* live and/or heat killed spores (f) were pre-incubated in bacterial supernatants as in (a) and treated with propidium iodide (f). The fraction of propidium iodide^+^ spores was quantified. (g) The fraction of fired spores post-supernatant incubation was quantified. (h) The concentration of spores 24 hours post-incubation was quantified. (i-k) *N. parisii* spores were pre incubated in supernatants denoted on the X-axis and then used to infect L1s for one hour. The number of spores per animal (i), the percentage of fired spores alongside a spore control of spores incubated overnight but not fed to animals (j), and the number of sporoplasms (k) are displayed. Data is from three independent replicates of 100 worms (b) or 20 worms each (c-e,i-k) and at least 50 spores each (c-d,f-j). Mean ± SD represented by horizontal bars. P-values determined via one-way ANOVA with post hoc. Significance defined as * p < 0.05, ** p < 0.01, *** p < 0.001, **** p < 0.0001.

Molecules could inactivate microsporidia spores by making them less viable, inducing germination, or by reducing spore numbers (Buczek et al. 2020; Han, Takvorian, and Weiss 2020). Propidium iodide (PI) is a dye that only stains non-viable cells. Spores treated with *P. lurida* MYb11 or *P. mendocina* MSPm1 supernatants did not display increased levels of PI staining compared to heat-killed spores, indicating that this was not the mechanism of action **(Figure 4f)**. We next tested if supernatant treatment induces premature spore firing *in vitro*. However, the fraction of fired spores was relatively similar under all treatment conditions making this scenario unlikely **(Figure 4g)**. Given that *P. lurida* MYb11 and *P. mendocina* MSPm1 supernatants reduce the number of spores in the nematode gut, we tested the possibility of *in vitro* spore destruction. Incubation of *N. parisii* spores in either supernatant resulted in a decreased concentration of spores *in vitro* **(Figure 4h)**. Lastly, we tested how a combination of *P. lurida* MYb11 and *P. mendocina* MSPm1 supernatants impact *N. parisii* infectivity. Incubation of *N. parisii* spores in the two supernatants resulted in a reduction in the number of spores in the gut without any alterations in spore firing **(Figure 4i,j)**. However, the number of intracellular sporoplasms was further reduced when both supernatants were present suggesting that these two bacteria produce different molecules **(Figure 4k)**.

### Massetolide E and F produced by *P. lurida* MYb11 inhibit *N. parisii* spores

*Pseudomonas* bacteria are known to produce a wide range of small molecules with antimicrobial activity (Nguyen et al. 2016). To isolate active compounds present within the supernatants of *P. lurida* MYb11 and *P. mendocina* MSPm1, we conducted activity guided purification followed by liquid chromatography-mass spectrometry. Measurement of purified compounds from *P. lurida* MYb11 by mass spectrometry identified two molecules, massetolide E and F **(Figure S8a,b)**. To test if massetolide E and F display anti-microsporidia activity, we incubated *N. parisii* in varying concentrations of these compounds. Although massetolide E and F do not significantly reduce the number of spores in the gut or alter spore firing, there is a reduction in the number of sporoplasms in intestinal cells **(Figure S8c,d,5a).** To test if these two compounds display additive or synergistic effects, 128 μg/ml of each compound was added to *N. parisii* spores. Under these treatment conditions, spores were also less infective as seen by the reduced number of sporoplasms, similar to when each compound was added in isolation at 256 μg/ml. These results indicate that massetolide E and F display anti-microporidia activity in an additive manner **(Figure 5a)**.

**Figure 5.**
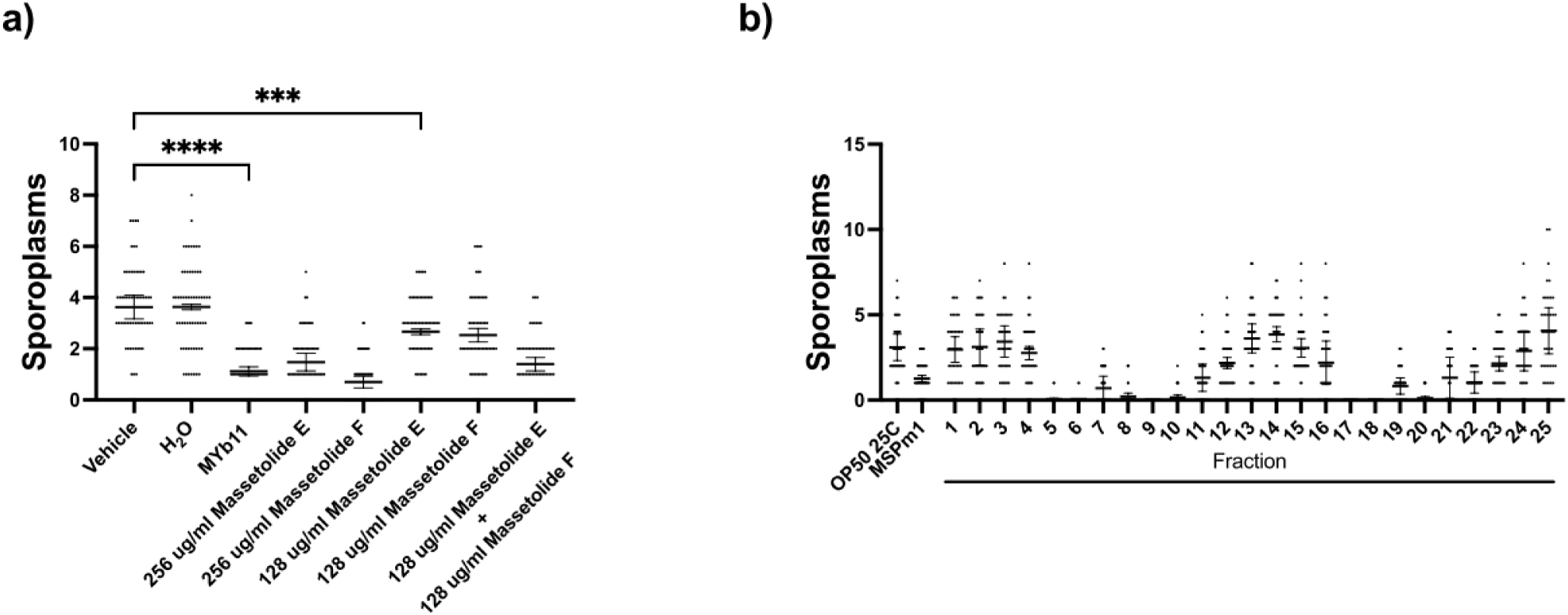
Massetolide E and F and two unknown molecules from *P. mendocina* MSPm1 exhibit anti-microsporidia activity. (a) Spores were incubated in either a vehicle control (0.5% DMSO), water, *P. lurida* MYb11 supernatant or massetolide E and/or F in various concentrations prior to L1 infection. The number of sporoplasms in L1 animals was determined. (b) Liquid chromatography of *P. mendocina* MSPm1 supernatant with a C18 column resulted in 25 fractions that were tested to identify those containing activity. The number of sporoplasms in L1 animals was quantified. Data is from two (b) or three (a) independent replicates of 20 worms each. Mean ± SD represented by horizontal bars. P-values determined via one-way ANOVA with post hoc. Significance defined as *** p < 0.001, **** p < 0.0001.

Cultured media from *P. mendocina* MSPm1 inhibits microsporidia spores in a similar fashion to *P. lurida* MYb11 however this strain is not predicted to produce massetolides **(Table S1)**. To identify what other anti-microsporidia compounds naturally exist, we performed activity-guided purification. The CombiFlash fractions showed activity in two different sets of fractions, suggesting the presence of two different molecules with anti-microsporidia activity. Two separate fraction intervals, 5-10 and 17-18, inhibit *N. parisii* spore infectivity as seen by a reduction in the number of sporoplasms when spores were incubated in with these fractions **(Figure 5b).**

### A diverse repertoire of *C. elegans*-associated *Pseudomonas* species display anti-microsporidia activity

*Pseudomonas* species (spp.) are frequently isolated from the native *C. elegans* habitat (Schulenburg and Félix 2017). We next tested whether other species of *Pseudomonas* also produce anti-microsporidia compounds. We screened a set of 53 *Pseudomonas* spp. associated with *C. elegans* by incubating *N. parisii* spores in cultured media from these bacterial isolates (Dirksen et al. 2016; Johnke, Dirksen, and Schulenburg 2020; Zimmermann et al. 2020). Synchronized L1 N2 animals were infected with pre-treated spores for 72 hours, and *N. parisii* infectivity was quantified by classifying nematodes as being lightly, moderately, or heavily infected. We observed that 64% (34/53) of the *Pseudomonas* isolates secrete molecules which reduce the proportion of heavily infected animals **(Figure S9a,b)**.

To determine the evolutionary relationship between species that displayed anti-microsporidia activity, we created a phylogenetic tree using *rpoD* sequences **(Figure 6a)** (Zimmermann et al. 2020; Lauritsen et al. 2021). We observed a distinct phylogenetic relationship between species with activity towards *N. parisii*. For example, all nine *P. lurida* isolates and all eight *P. canadensis* isolates we tested significantly inhibit infection. Conversely, only one out of seven *P. fluorescens* isolates tested has significant activity towards *N. parisii*. In total, we observed eight named species and at least 13 phylogenetically distinct groups of *Pseudomonas* with activity against *N. parisii*. To investigate the arsenal of secreted compounds produced by *C. elegans*-associated *Pseudomonas* species, we used antismash analysis to identify candidate bacterial gene clusters (BGC) from 21 genomes, including six we sequenced for this study (Blin et al. 2023). All seven *P. lurida* isolates’ genomes are predicted with high confidence to encode for viscosin, as was the one *P. canadensis* genome analyzed, MYb395. No other genomes are predicted to confidently make viscosin, though they are predicted to make a diversity of natural products, albeit with lower confidence **(Table S1)**. Together these results suggest that a diverse group of *C. elegans*-associated *Pseudomonas* species commonly produce a variety of molecules which inhibit *N. parisii* infection.

**Figure 6.**
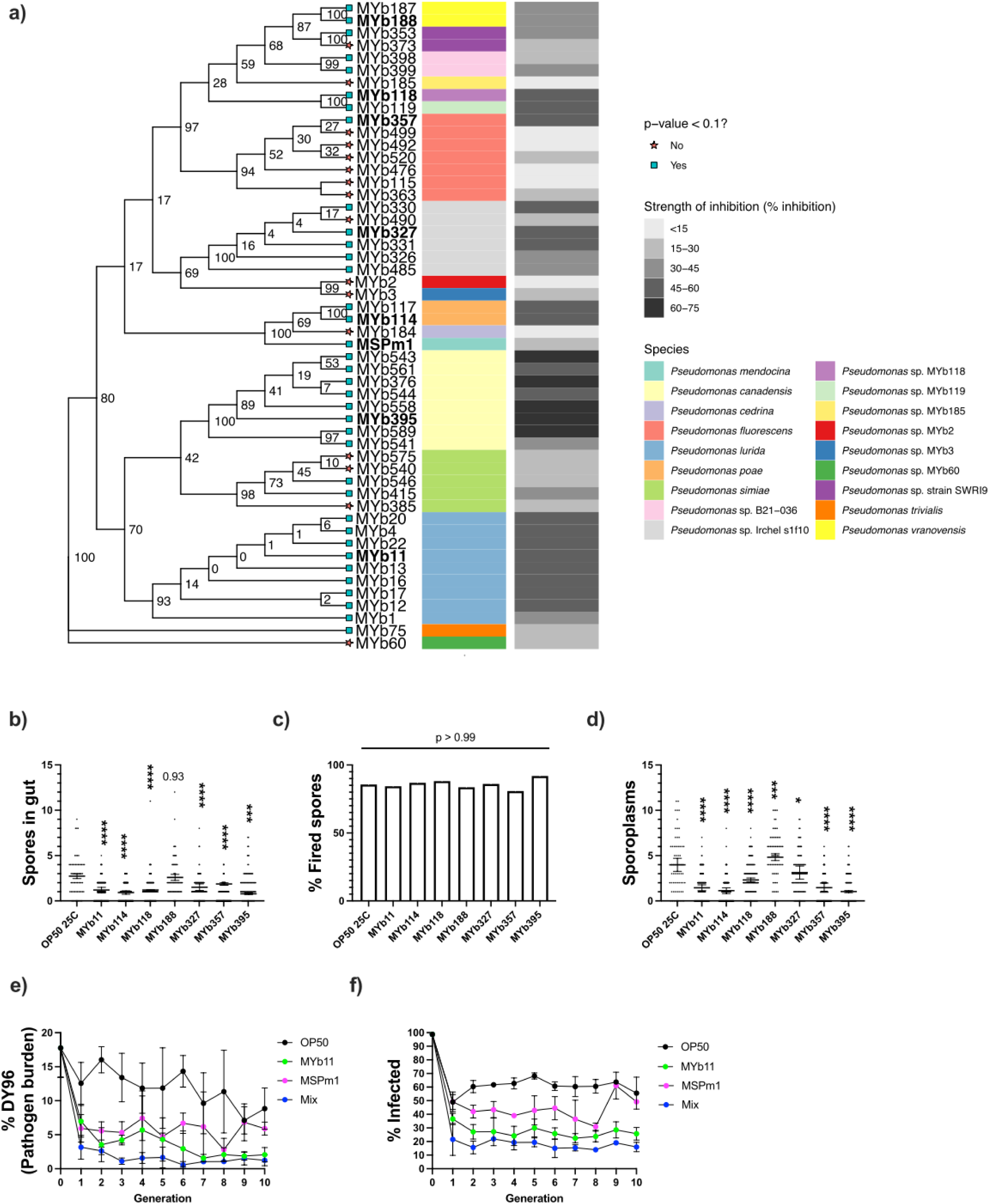
Many *Pseudomonas* species inhibit *N. parisii* spores and infection progression over multiple generations. (a) Phylogenetic tree demonstrating the evolutionary relatedness between *Pseudomonas* isolates using *rpoD* sequences. The isolates used in downstream experiments are bolded. The different *Pseudomonas* species are indicated using a color gradient. The strength of inhibition exhibited by the cultured media from each strain are indicated in varying shades of grey to black where black indicates the highest strength of inhibition. Blue squares indicate significant inhibitory effects of p < 0.1. (b-d) L1 animals were pulse infected for one hour with *N. parisii* spores incubated in supernatants of bacterial strains depicted on the X-axis. The number of spores (b), the fraction of fired spores (c), and the number of intracellular sporoplasms (d) were quantified. (e-f) L1 animals were infected for 72 hours and adults were propagated for 10 generations on various lawns of bacteria. Mix condition contains MYb11, MSPm1 and the 6 tested isolates from b-d in equal proportion. The pathogen burden (e) and the fraction of population infected are depicted (f). Data is from three independent replicates of at least 20 (b-e) or 100 (f) worms each and at least 50 spores (b,c). Mean ± SD represented by horizontal bars. P-values determined via one-way ANOVA with post hoc. Significance defined as * p < 0.05, ** p < 0.01, *** p < 0.001, **** p < 0.0001 and is relative to *E. coli* OP50 25°C.

In the wild, *C. elegans* are likely to be exposed to both *N. parisii* and *Pseudomonas* isolates, and molecules produced by these bacteria could act on spores as well as intracellular stages after *N. parisii* invasion. To determine mechanisms of how other *Pseudomonas* species inhibit *N. parisii*, we first incubated spores with conditioned media from six phylogenetically distinct *Pseudomonas* isolates and exposed L1 worms to these treated spores. The incubation of *N. parisii* in 5 of the 6 *Pseudomonas* isolates tested resulted in fewer spores and sporoplasms in the nematode gut and no difference in the fraction of fired spores **(Figure 6b-d).** We then pulse-infected L1 animals and placed them onto individual *Pseudomonas* lawns for the next 71 hours. When animals were pulse infected, MYb11, MYb188, and MYb327 lawns resulted in improved host fitness relative to *E. coli* OP50 and we observed a small but significant decrease in pathogen burden **(Figure S10a-c)**. In addition, we set up continuous infections on lawns of the different *Pseudomonas* species, such that parasite invasion and proliferation are taking place in the presence of these bacteria. When animals were continuously infected, all *Pseudomonas* spp. resulted in improved host fitness with decreased pathogen burden in all but MYb118, MYb188, and MYb327 exposed animals **(Figure S10d-f)**.

### *Pseudomonas* species provide strong protection against *N. parisii* infection over multiple generations

Native microbiomes have the potential to limit infectious disease (Libertucci and Young 2019; Gupta, Singh, and Mani 2022). In the wild, *C. elegans* is often exposed to a variety of different bacteria (Samuel et al. 2016; Dirksen et al. 2020; 2016; Zimmermann et al. 2020). As individually tested *Pseudomonas* species could reduce microsporidia infection, we asked if combining *Pseudomonas* species together may result in stronger effects. First, we tested if continuous infection of animals on lawns of *P. lurida* MYb11, *P. mendocina* MSPm1, or a combination of the two would similarly impact *N. parisii* infectivity. Host fitness was significantly improved when grown on single or mixed lawns relative to when grown on *E. coli* OP50 **(Figure S11a)**. Growth on either *P. lurida* MYb11 or *P. mendocina* MSPm1 resulted in decreased parasite burden and combining these two strains did not result in a stronger protective effect **(Figure S11b)**.

We then asked how a larger combination of bacterial species may impact *N. parisii* infection over time. To do this, we continuously infected L1 animals for 72 hours and picked 10 gravid animals from the same source plate onto lawns seeded with either *E. coli* OP50, *P. lurida* MYb11, *P. mendocina* MSPm1 or a mix of eight *Pseudomonas* species (MYb11, MSPm1, MYb114, MYb118, MYb188, MYb327, MYb357 and MYb395). Animals were propagated for 10 generations, and the pathogen burden and population infectivity were quantified at every generation. Although animals grown on *E. coli* OP50 showed a slow decline in pathogen burden, the population remained heavily infected **(Figure 6e,f)**. Animals grown on *P. lurida* MYb11, *P. mendocina* MSPm1 or the 8 species mix displayed faster decline in pathogen burdens and population infectivity. Although the pathogen burden continued to decline past generation 8 for *P. lurida* MYb11 and the mix conditions, a sharp increase was observed for those grown on *P. mendocina* MSPm1. Both *P. lurida* MYb11 and mix conditions resulted in continually low levels of *N. parisii* infection, and the mix condition resulted in a lower proportion of the population producing spores **(Figure 6e,f)**.

## Discussion

To understand how bacterial members of the native *C. elegans* microbiome impact nematodes infected with *N. parisii,* we utilized the recently established CeMbio collection of bacteria. Here, we demonstrate that bacterial species can alter *N. parisii* infection in multiple ways. First, we show that nutrient limitation can restrict *N. parisii* of the required host nutrients to fuel its growth. Second, *Pseudomonas* bacteria make molecules that target dormant spores and growth on these bacteria reduces infection within populations for many generations. Altogether, this work demonstrates that the host bacterial microbiome can exert large impacts on microsporidia infection. Although the panel of bacteria we assayed covers a large extent of the known diversity of the *C. elegans* microbiome, other species and combinations of bacteria may also have an impact on microsporidia infection (F. Zhang et al. 2017; Dirksen et al. 2020).

Microsporidia infection can be influenced by host diet (Franchet et al. 2019; Porrini et al. 2011; Willis and Reinke 2022). We observed that worms grown on *S. multivorum* BIGb0170 or *C. scopthalmum* JUb44 cause an increase of microsporidia invasion, which likely occurs by increased pumping in the nematodes due to a poor quality food source (Avery and Horvitz 1990; Dirksen et al. 2020). Despite this initial increase in pathogen numbers, the ability to sporulate is delayed on these bacteria, and metabolomics revealed that these diets caused a disruption to host unsaturated lipid levels. Microsporidia infection in multiple animals has been shown to result in decreased fatty acid levels, including reduced linoleic acid in the gypsy moth (Franchet et al. 2019; Su et al. 2023; Ning et al. 2019; Hoch et al. 2002). Fatty acid supplementation of either palmitic or oleic acid in *Drosophila melanogaster* resulted in increased proliferation of the microsporidian *Tubulinosema ratisbonensis* (Franchet et al. 2019). Protection against pathogenic bacteria by beneficial bacteria through alterations of host sphingolipid levels has also been observed in *C. elegans* (Peters et al. 2024). We found that the polyunsaturated fat, linoleic acid, can restore *N. parisii* infectivity in nematodes grown on *S. multivorum* BIGb0170 and depletion of linoleic acid through the loss of the *C. elegans* desaturase FAT-2 delays *N. parisii* sporulation, suggesting an important requirement of polyunsaturated fatty acids for microsporidia infection.

Bacteria can inhibit microsporidia infection either through activating host immunity or by producing molecules which directly act on microsporidia. Infection with microsporidia induces a transcriptional program called the intracellular pathogen response, and only about 7% of bacteria associated with *C. elegans* activate this immune response (González and Félix 2024; Reddy et al. 2017; Wan, Troemel, and Reinke 2022). This includes *C. piscis* BIGb0172, for which we observe a slight decrease in infection. This suggests that immune activation by the microbiome may play a minor role in protection against microsporidia infection of *C. elegans*. In contrast, we observed that 2 out of the 11 CeMbio species tested produce molecules that cause an ∼2-4-fold reduction in invasion. Both of these inhibitory bacteria are *Pseudomonas* species and 65% of *Pseudomonas* isolates we tested make molecules that have activity against *N. parisii* spores. About 22% of all isolated *C. elegans*-associated bacteria are *Pseudomonas*, which suggests an estimate of about 14% of the bacteria *C. elegans* encounters in the wild provide protection against *N. parisii* spores (Dirksen et al. 2016; 2020; Samuel et al. 2016). We also observed that *Pseudomonas* isolates can inhibit *N. parisii* proliferation, suggesting that compounds produced by these bacteria can act on intracellular stages of microsporidia. Although many *Pseudomonas* species are pathogenic against *C. elegans*, of the eight isolates with activity against microsporidia, only two caused a reduction in *C. elegans* fitness, suggesting that many *Pseudomonas* species could have a beneficial effect on *C. elegans* in the wild. These *Pseudomonas* bacteria also produce vitamin B12, which provides tolerance to *N. parisii* infection (Zimmermann et al. 2020; Willis et al. 2023). As bacteria seldom exist in isolation, we mimicked natural conditions by growing infected nematodes on lawns containing a mixture of bacteria and showed that a mixture of *Pseudomonas* could provide robust protection against *N. parisii* for multiple generations.

*Pseudomonas* species produce many molecules with antimicrobial activities (Nguyen et al. 2016; Dirksen et al. 2016). We identified that *P. lurida* MYb11 produces massetolides E and F, which are cyclic lipopeptides related to viscosin that have been demonstrated to have wide antimicrobial activity with effects on viruses, protozoa, oomycetes and bacteria (de Bruijn et al. 2007; De Souza et al. 2003; Gerard et al. 1997; Kissoyan et al. 2019; Raaijmakers, de Bruijn, and de Kock 2006; Geudens and Martins 2018). *P. lurida* MYb11 has been previously reported to produce massetolide E, which was shown to display potent antibacterial activity against the gram-positive bacterial pathogen *B. thuringiensis*. The mode of action of many cyclic lipopeptides occurs through disruption of target cell membrane integrity, resulting in cell death (Schneider et al. 2014; van de Mortel et al. 2009). This process has been well documented in oomycetes such as massetolide A inducing zoosporicidal activity against the late blight plant pathogen *Phytophtora infestans* (van de Mortel et al. 2009). We demonstrate that massetolide E and F impact *N. parisii* spores through a reduction in the number of spores, which is an inhibitory mechanism that has been observed with other molecules such as porphyrins (Buczek et al. 2020). Several species of gram-positive *Bacilli* bacteria have been shown to produce antimicrobial activity towards microsporidia, suggesting that bacterial inhibition of microsporidia might be common (Porrini et al. 2010; Mossallam, Amer, and Diab 2014; X. Zhang et al. 2022).

Microsporidia are widespread parasites of agriculturally important animals and are a threat to food security, but there are limited approaches to treating or preventing these infections. The most commonly used anti-microsporidia drugs suffer from either host toxicity or limited activity against some species (Han and Weiss 2018). *C. elegans* has been used to identify novel microsporidia inhibitors from small molecule libraries and we now show that this model organism can also be used to identify bacteria that produce anti-microsporidia compounds (Murareanu et al. 2022; Qingyuan Huang et al. 2023). There is an interest in using bacteria to combat infections, and probiotics have been used in honey bees to reduce microsporidia infections, though with mostly modest results (Alberoni et al. 2016). The demonstration that *Pseudomonas* bacteria produce a diversity of molecules with activities against microsporidia provides a potential starting point to harness these bacteria to control microsporidia infections.

## Acknowledgments

We are grateful to Qingyuan Huang, Claire Turke, Angcy Xiao and Winnie Zhao for providing helpful comments on the manuscript. We thank Hinrich Schulenburg for providing us the CeMbio strains and the *Pseudomonas* isolates. We thank Johannes Zimmermann for performing initial bioinformatic characterization of the metabolic capabilities of BIGb0170 and JUb44. Additional *C. elegans* strains were provided by the *Caenorhabditis* Genetics Center, which is funded by the National Institutes of Health (NIH) Office of Research Infrastructure Programs Grant P40 OD010440. Schematics were created using BioRender.com.

## Data availability

All experimental data is presented in Data S1 and all sequencing data is deposited in NCBI.

## Materials and Methods

### Strain Maintenance

*C. elegans* strains were maintained at 21°C on nematode growth medium (NGM) plates seeded with 10x *Escherichia coli* OP50-1. For all infection assays, 15 L4 animals were picked onto a 10-cm seeded NGM plate. 4 days later, heavily populated plates were bleach synchronized as previously described (Tamim El Jarkass et al. 2022). Embryos were hatched overnight in 5 ml of M9 at 21°C. L1 animals were used no later than 20 hours post bleaching. For a list of strains utilized in this study, refer to **Table S2**.

For 72 hours of growth on CeMbio strains, 1,000 synchronized L1 animals were grown on 6-cm NGM plates seeded with individual microbiome strains and incubated at 20°C. 24 **(Figure S1a-d)** or 72 hours later **(Figure 1)**, animals were washed off with M9 + 0.1% tween-20 into individual microcentrifuge tubes. These animals were then washed twice more with M9 + 0.1% tween-20 to remove residual bacteria from the supernatant.

### Bacterial growth and Maintenance

*E. coli* OP50-1, CeMbio strains and *Comamonas aquatica* DA1877 were struck out onto LB agar and incubated at 25°C for 24 hours or 48 hours (BIGb0170, BIGb0172, MYb1, MYb115, and MYb185). Colonies were kept for a maximum of two weeks at 4°C. JUb134 was excluded as it did not grow at a similar rate to other strains in overnight cultures. Individual colonies were then picked into 5 ml of LB or 2x Tryptic soy broth (TSB) (MYb357) and cultured overnight at 25°C for 16-18 hours at 220 RPM. Cultures were then adjusted to an OD_600_ of 1.0 using a Microspek^TM^ DSM3 cell density meter. Cultures were either diluted in LB or concentrated via centrifugation at 7,197 rcf for 5 minutes. 180 μl of culture was used to seed a 6-cm NGM plate and 450 μl (Metabolomics) or 1 ml (10 generation experiments) for a 10-cm NGM plate. For experiments involving *E. coli* OP50 supplementation, 6-cm NGM plates were seeded with 180 μl in a 1:1 ratio of *E. coli* to JUb44 or BIGb0170 (90 μl each). Seeded plates were incubated at 25°C for 24 hours (or 4 days when 1 ml of culture was used) and stored at 4°C until use. Freshly seeded plates were used for every experimental replicate except in the case of multigenerational experiments (see below). *E. coli* OP50-1 at 37°C represents a culture grown at 37°C and saturated 10x. This was also used at an OD_600_ of 1.0, and was adjusted with LB. This served as a control for *E. coli* OP50 cultured at 25°C.

### Dietary supplementation

Fatty acids were supplemented post-autoclave to standard 6-cm NGM plates at 0.8 mM as previously described (Deline, Vrablik, and Watts 2013). Briefly, saturated fatty acids (palmitic and stearic acid) were resuspended in 100% ethanol and heated at 65°C to dissolve. NGM was supplemented, prior to autoclaving, with tergitol at a final concentration of 0.1% for ethanol, palmitic and stearic acid supplemented plates.

Ethanol plates were prepared at a final concentration of 0.5%. NGM flasks were weighed prior to autoclaving and brought back to their starting weight with autoclaved water once sterilized. Plates were stored at 4°C until use. Seeded plates were incubated with desired bacterial species at 25°C for 24 hours to promote bacterial uptake of fatty acids prior to the addition of *C. elegans*. Palmitic acid [Sigma-Aldrich P0500], stearic acid [Sigma-Aldrich 175366], oleic acid [Sigma-Aldrich O1008], linoleic acid [Sigma-Aldrich L1376] and tergitol [Millipore Sigma-NP40S] were utilized in this study.

### L1 pulse infection

For 1 hour pulse, 71-hour chase experiments either 25,000 bleach-synchronized N2 L1 animals were infected with a high dose of *N. parisii* (ERTm1) spores **(Figure 3)** or 6,000 bleach synchronized N2 L1 animals were infected with a medium dose of *N. parisii* spores **(Figure S5, S6b-c).** In both circumstances, 10 μl 10X *E. coli* OP50-1, or a mock treatment of an equivalent volume of M9 instead of spores (uninfected) was added. Worms, *N. parisii* spores/M9, and *E. coli* were placed in a microcentrifuge tube and mixed via pipetting prior to plating on an unseeded 6-cm NGM plate. After one hour, worms were washed off plates using 700 μl of M9 + 0.1% Tween-20 and placed into a microcentrifuge tube. Animals were then washed twice in M9 + 0.1% Tween-20 to remove residual spores in the supernatant prior to evenly splitting the worm population onto seeded plates of choice.

For L1 pulse infections with spores incubated in supernatants **(Figure 4b-g, i-k** ,**6b-d, S7 and S9b)**, fractions **(Figure 5b)**, or compounds **(Figure 5a and S8)**, 1,000 bleach synchronized N2 L1 animals and 10 μl 10X *E. coli* OP50-1 were added to the microcentrifuge tube containing *N. parisii* spores and supernatant, fraction or compound and plated onto an unseeded 6-cm NGM plate.

For a list of spore doses used in this study, refer to **Table S3.**

### Adult pulse infection

72-hour old animals were infected by adding a medium dose of *N. parisii* spores and 10 μl of saturated 10x OP50-1 into the microcentrifuge tube containing the adult animals. This mixture was gently pipetted up and down prior to placing it onto an unseeded 6-cm NGM for one hour at 20°C. Animals were then washed off with M9 + 0.1% tween-20. Two additional M9 + 0.1% tween-20 washes were performed to remove residual spores that have not been ingested. A fraction of these animals was set aside for fixation (representing 1hpi) and the rest were placed onto a 6-cm NGM plate seeded with the corresponding bacteria on which they were grown. 24, 48 or 72 hours later, animals were washed off with M9 + 0.1% tween-20 and fixed in acetone for downstream FISH and direct yellow 96 (DY96) staining.

### Quantifying pharyngeal pumping

72-hour old animals were picked onto 6-cm *E. coli* OP50 seeded NGM plates immediately prior to measurement. Pumping was measured for one minute per animal using an Axio Zoom V.16 (Zeiss).

### Lipidomics

2,500 N2 L1 animals were placed onto each of two 10-cm plates seeded with the desired bacterial species (450 µl) for 72 hours at 20°C. All animals or bacteria were then washed with 1 ml of M9 into a microcentrifuge tube. For samples containing nematodes, animals were washed to remove residual bacteria from the sample. All samples were then flash frozen in liquid nitrogen and stored at -80°C until use in extraction.

Lipids were extracted from pelleted and frozen animals using Bligh-Dyer extraction (Bligh and Dyer 1959) in a bead mill homogenizer. Samples were first mixed via vortexing and pipette agitation and resuspended in ice-cold methanol. One volume of methanol-sample mixture was transferred to one volume of ice-cold chloroform in beadmill homogenizer tubes. Then, accounting for water content in the initial sample, water was added to bring the final water content to 0.9 volumes. Following homogenization and phase-separation by centrifugation, lower organic layers were collected for lipidomics, and dried in a vacuum evaporator.

The dried organic layer was resuspended in 50µL of 1:1 AcN:IPA (v/v). Reconstituted extracts were analyzed with a Vanquish dual pump liquid chromatography system coupled to an Orbitrap ID-X (Thermo Fisher Scientific) using a H-ESI source in positive mode. All samples were injected at 2 μL and analytes separated with 30 minute reversed-phase chromatography (Accucore C30 column; 2.6 μm, 2.1mm × 150mm; 27826-152130; Thermo) with an Accucore guard cartridge (2.6 μm, 2.1 mm × 10 mm, 27826-012105, Thermo). Mobile phase A consisted of 60% LC/MS grade acetonitrile (A955, Fisher). Mobile phase B consisted of 90% LC/MS grade isopropanol (A461, Fisher) and 8% LC/MS grade acetonitrile. Both mobile phases contained 10mM ammonium formate (70221, Sigma) and 0.1% LC/MS grade formic acid (A11710X1-AMP, Fisher). Column temperature was kept at 50 °C, flow rate was held at 0.4 mL/min, and the chromatography gradient was as follows: 0-1 min held at 25% B,1-3 min from 25% B to 40% B, 3-19 min from 40% B to 75% B, 19-20.5 min from 75% B to 90% B, 20.5-28 min from 90% B to 95% B, 28-28.1 min from 95% B to 100% B, and 28.1-30 min held at 100% B. A 30 minute wash gradient was run for every column injection to clean up the column and re-equilibrate solvent conditions that went as follows: 0-10 min held at 100% B and 0.2 mL/min, 10-15 min from 100% B to 50% B and held at 0.2 mL/min, 15-20 min held at 50% B and 0.2 mL/min, 20-25 min from 50% B to 25% B and held at 0.2 mL/min, 25-26 min held at 25% B and ramped from 0.2 mL/min to 0.4 mL/min, and 26-30 min held at 25% B and 0.4 mL/min. Mass spectrometer parameters were: source voltage 3250V, sheath gas 40, aux gas 10, sweep gas 1, ion transfer tube temperature 300°C, and vaporizer temperature 275°C. Full scan data was collected on experimental replicates using the orbitrap with scan range of 200-1700 m/z at a resolution of 240,000 FWHM. On pooled samples for lipid identification, primary fragmentation (MS2) was induced in the orbitrap with assisted HCD collision energies at 15, 30, 45, 75, 110%, CID collision energy was fixed at 35%, and resolution was at 15,000. Secondary fragmentation (MS3) was induced in the ion trap with rapid scan rate and CID collision energy fixed at 35% for 3 scans. LipidSearch (v 5.0, Thermo) was used for lipid annotation and used to generate a mass list for peak picking and integration of experimental replicates in Compound Discoverer (v 3.3).

Data was analyzed with in-house scripts in R and heatmaps were generated via the pheatmap R package.

### Sample fixation and staining

Samples were washed off plates with 1 ml of M9 + 0.1% Tween-20 into a microcentrifuge tube. They were then washed once more before adding 700 μl of acetone to worm pellets. DY96, a chitin binding dye, was used to assess parasite burden (microsporidia) and worm embryos. 500 μl of DY96 solution (1 x PBST, 0.1% SDS, 20 μg/ml DY96) was added to worm pellets that had been fixed in acetone. Samples were left to rock in the dark for 30 minutes at room temperature and then centrifuged to remove the dye. Worms were then resuspended in 20 μl of EverBrite Mounting Medium (Biotium) and 10 μl was mounted onto glass slides for imaging. To observe early intracellular infection events (sporoplasms), the MicroB FISH probe (5’-ctctcggcactccttcctg-3’) conjugated to Cal Fluor 610 (LGC Biosearch Technologies) was used to bind the 18S rRNA of *N. parisii.* Animals fixed in acetone were washed twice in 1 ml PBST, followed by a single 1ml wash in Hybridization buffer (0.01% SDS, 900 mM NaCl, 20 mM TRIS pH 8.0). Samples were then incubated overnight at 46°C in the dark with 5 ng/μl of the MicroB FISH probe in 100 μl of Hybridization buffer. Samples were then washed in 1 ml of wash buffer (Hybridization buffer + 5mM EDTA), and incubated in 500 μl of wash buffer for 30 minutes at 46°C. To visualize intracellular stages (sporoplasms and meronts) alongside spores, the final incubation was replaced with 500 μl of DY96 solution for 30 minutes at room temperature. The supernatant was then removed, and samples were resuspended in 20 μl of EverBrite Mounting Medium (Biotium).

### Spore Firing Assays

Animals stained with FISH and DY96 were used to determine the fraction of fired spores. FISH^+^ DY96^+^ spores represent unfired spores, whereas FISH^-^DY96^+^ spores represent fired spores. FISH^+^DY96^-^ events represent intracellular sporoplasms. Percentage of fired spores is defined as the proportion of fired spores over the total number of fired and unfired spores within a population.

### Microscopy and Image quantification

All imaging was performed using an Axio Imager.M2 (Zeiss), except for quantification of *Pacdh-1::*GFP in **(Figure S6a)**, which was imaged using an Axio Zoom V.16 (Zeiss) at a magnification of 45.5x. Images were captured via Zen software and quantified under identical exposure times per experiment. Gravidity is defined as the presence of at least one embryo per worm, and animals were considered infected if clumps of newly formed spores (48-72 hpi) were visible in the body of animals as seen by DY96. FISH-stained animals were considered infected if at least one sporoplasm was visible in intestinal cells.

To quantify fluorescence within animals (pathogen burden), regions of interest were used to outline every individual worm from anterior to posterior. Individual worm fluorescence from DY96 or FISH staining were subjected to the “threshold” followed by “measure” tools in FIJI (Schindelin et al. 2012; Willis, Jarkass, and Reinke 2022).

### *In vitro* spore incubation

Overnight cultures of bacterial strains were grown as described above. Cultures were then centrifuged at 7,197 rcf for 5 minutes, and 100 μl of supernatant, massetolide or fractions were placed into a microcentrifuge tube with a medium dose of *N. parisii* spores at 20°C for 21-24 hours. The entire volume was used for downstream experiments. For continuous infections using wild *Pseudomonas* isolates, a low dose of *N. parisii* spores was used **(Figure S9b)**.

Massetolide E and F were resuspended in 100% DMSO to a final concentration of 5 mg/ml and stored at -20°C. 256 μg/ml and 128 μg/ml concentrations were generated by adding 5.12 and 2.56 μl of 5 mg/ml stocks respectively in a final volume of 100 μl of nuclease free water. Vehicle controls contained 5μl of 100% DMSO in 95 μl of nuclease free water. H_2_0 control represents spores incubated in 100 μl of nuclease free water.

To wash spores incubated in bacterial supernatant, samples were centrifuged at 12,000 x g for 1 minute. Spores were washed twice in 500 μl of autoclaved water and resuspended in the same volume as the unwashed samples.

### 72-hour continuous infections

1,000 bleach synchronized L1 animals and 400 μl of 10x *E. coli* OP50-1 were added to the microcentrifuge tube containing the supernatant-spore incubations **(Figure 4b, S7)**. Samples were then plated onto unseeded 6-cm NGM plates for 72 hours at 20°C. Animals were washed off with 1 ml of M9 + 0.1% Tween-20 and fixed in acetone. When using seeded plates **(Figure S10d-f, S11),** 1,000 bleach synchronized L1 animals and a high dose of *N. parisii* spores was added to a final volume of 200 μl in M9 and plated onto a seeded 6-cm NGM plate.

### Propidium Iodide staining

Propidium iodide (P4170-Sigma Aldrich) was added to samples at a final concentration of 1μg/ml after overnight incubation in bacterial supernatants. Heat killed spores were generated as a positive control for PI staining by incubating spores in 100 μl of LB at 65°C for 10 minutes.

### Quantifying spore concentrations

4 μl of Calcofluor white (CFW) was added to pre-incubated spores and mixed via pipetting. 4 μl of this sample was then mounted onto a Cell-Vu slide (Millenium Sciences DRM600). The number of spores in 10 squares of the grid was measured three independent times and averaged to represent a single biological replicate. Spore concentration was calculated as described by the manufacturer (# of spores in 10 grids / 2 = million spores/ml). Spore samples were blinded prior to quantification.

### Fermentation and purification of anti-microsporidia compounds

*P. lurida* MYb11 and *P. mendocina* MSPm1 were grown in TSB medium. 50 ml of inoculum in TSB was used for inoculating a 1-liter culture. Both the stains were grown at 25°C at 220 rpm. After 48h, the supernatant was separated from the cell by centrifugation at 6,000 rpm for 20 min. The cell-free supernatant was added to 6-8-% (w/v) of activated Diaion HP-20 (Sigma) resin. This mixture was allowed to mix for 2-3h. The resins were separated and washed with distilled water. The compounds that were bound to the resin were eluted with 100% methanol. The solvent was evaporated using a rotary evaporator (RotaVap, Heidolph). The extract was reconstituted in Milli-Q water. The extract was loaded onto a Sephadex LH 20 column and eluted using 50% methanol in isocratic mode. The fractions were collected and concentrated using GENEVAC evaporator. The fractions were reconstituted using Milli-Q water and the activities were measured as described above. The active fractions were pooled and loaded on reverse-phase CombiFlash ISCO column (RediSep Gold Rf C18, Teledyne) C. The compounds were eluted with acetonitrile (ACN) and water with 0.1% formic acid using a gradient of 5% to 95% ACN. The fractions were concentrated and the activities were assessed. The presence of massetolide E and massetolide F from the *P. lurida* MYb11 strain was detected by using mass spectrometry in the CombiFlash fractions. The active fractions were dried and then applied to an HPLC C28 Analytical column (Sunniest RP-AquA C28 4.6X 100 mm, 5µm) using water and eluted with a gradient of ACN. Both solvents contained 0.1% formic acid. The fractions were collected and the activity was assessed. The active peaks eluted after 95% of ACN. The pure compounds, massetolide E and massetolide F, were lyophilized to generate a white powder.

In the case of *P. mendocina* MSPm1, the CombiFlash fractions indicated the presence of two different compounds exhibiting anti-microsporidia activity. The two compounds were different in terms of their polarity. The polar compounds eluted in earlier fractions of CombiFlash and the non-polar compounds eluted later. The yield of these compounds was very low. To mitigate this, we scaled up the fermentation and inoculated six liters of TSB with *P. mendocina* MSPm1. After 48h of incubation, we prepared and processed the extract as previously described. The polar fractions with activity were pooled and injected into HPLC on a C28 Analytical column (Sunniest RP-AquA C28 4.6X 100 mm, 5µm) using water and eluted with a gradient of ACN with 0.1% Formic acid. We collected seven peaks separately but did not get any significant activities. To overcome this, we decided to collect the individual peaks in multiple injections to get enough compound (at least 1mg each) to test the activity and identify the active compound. The non-polar fractions also showed decreased antibacterial activity. After several attempts of HPLC (analytical column X Select® C18 4.6X 100 mm, 5µm) injections, we did not get enough compound to test the mass.

### Mass Spectrometry

The mass of massetolide E and massetolide F were analyzed using HR-ESI-MS recorded with Infinity II LC System (Agilent Technologies) coupled with a qTOF 6550 mass detector in positive ion mode. The compounds were dissolved in DMSO and ran on qTOF using an Agilent C8 column (Eclipse XDB-C8 2.1X100mm, 3.5 µm) using water and acetonitrile with 0.1% formic acid as solvents.

### *rpoD* gene amplification and sequencing

1 ml of overnight culture was used for genome extraction of most of the *Pseudomonas* isolates using a bacterial genomic DNA extraction miniprep kit (Bio Basic-BS423) following the manufacturer’s instructions except for modifying the incubation time of buffer-treated bacteria to 2 hours at 56°C. Centrifugation steps were modified to 3 minutes at 21 rcf,130 rcf or 9,000 rcf. DNA was eluted with 35 µl nuclease-free H_2_O. Extracts were stored at -20°C. In the case of MYb476, MYb492, MYb520, MYb522, MYb543, and MYb544, single colonies of each strain were resuspended in 50 µl Milli-Q water in 1.7 ml Eppendorf tubes. The tubes were immersed in liquid nitrogen and then placed in a hot plate (70°C) three times. Lysates were stored at -20°C.

*rpoD* was then amplified using psEG30F (5’-ATYGAAATCGCCAARCG-3’) and psEG790R (5’-CGGTTGATKTCCTTGA-3’) and KAPA2G Fast Hotstart Readymix (Roche-07960956001) to generate a 736 bp product as previously described (Girard et al. 2020). 1 µl of 50 ng/µl whole genomic DNA or 1-2 µl of lysate was used as the PCR template. Amplicons were then PCR purified using a Monarch® PCR & DNA cleanup kit (NEB-T1030S) once a band of the correct size was visualized on a 1% Agarose gel. Sanger sequencing using the psEG790R primer was performed and high quality ∼600 bp amplicons were used to generate the phylogenetic tree (see below). *rpoD* sequences for MYb114 (NZ_PCQN01000019.1), MYb117 (NZ_PCQL01000021.1), MYb184 (NZ_PCQE01000032.1), and MSPm1 (NZ_CP059139.1) were acquired from NCBI. Sequences of *rpoD* generated in this study were deposited in NCBI with the accession numbers PP861230-PP861277.

### Species designation

*rpoD* sequences from each isolate were queried using BLAST (basic local alignment search tool https://blast.ncbi.nlm.nih.gov/Blast.cgi). Isolates were assigned a species group based on the following parameters: In the case where both NCBI and BLAST displayed different species predictions, those from NCBI were used. If NCBI did not assign a species name (strain only) then the species designation based on BLAST results with a >98% match to a whole genome sequence was used. If no match to a whole genome sequence was found via BLAST but NCBI provided a strain name, the species was determined via a BLAST match of >98% to the *rpoD* sequence. For MYb60 there was a 100% match to a single *P. fluorescens rpoD* sequence, but this isolate clustered separately from the other *P. fluorescens* isolates during phylogenetic analysis (below) and was therefore labelled as *P. sp*. MYb60.

### Phylogenetic analysis

ModelTest-NG v0.1.7 was run on the fasta alignment to estimate the best RAxML model. AIC and AICc statistical criteria suggested the general time reversible model (GTR+G4) (Darriba et al. 2020). Therefore, the phylogenetic tree was constructed with RAxML-NG v. 1.2.1 using the GTR+G4 model and the bootstrap values were calculated based on 1000 replications (Kozlov et al. 2019). Finally, the tree was rooted with MYb60 and plotted as a dendrogram using R packages ape v5.7.1 (Paradis and Schliep 2019), ggtree v3.10.0 (Yu 2022; Xu et al. 2022; Yu 2020; Yu et al. 2018; 2017), ggplot2 v3.4.4 (Wickham 2016), ggnewscale v0.4.9 (Campitelli 2024), ggtreeExtra v1.12.0 (Yu 2022; Xu et al. 2021), ggstar v1.0.4 (Xu 2022), and RColorBrewer v1.1.3 (Neuwirth 2022).

### Whole genome sequencing

Previously isolated genomic DNA from MYb114, MYb118, MYb188, MYb327, MYb357 and MYb395 were submitted for whole genome sequencing at Plasmidsaurus (https://www.plasmidsaurus.com/). Bacterial genome sequencing was performed by Plasmidsaurus using Oxford Nanopore Technology with custom analysis and annotation. Sequences were deposited in NCBI with the accession numbers pending.

### Antismash analysis

Genome assemblies were uploaded onto antismash (https://antismash.secondarymetabolites.org/#!/start) and ran using default parameters (Blin et al. 2023).

### Multigeneration continuous infections

Prior to the start of the experiment, unseeded 10-cm NGM plates were seeded with 1 ml of OD 1.0 *E. coli* OP50, or MYb11 or MSPm1 or an even mixture of MYb11, MSPm1, MYb114, MYb118, MYb188, MYb327, MYb357 and MYb395 – “Mix” (250 μl of each). Plates were incubated at 25°C for 4 days to promote lawn growth and stored at 4°C until use. The plates used to grow generation 1-7 were seeded with cultures originating from a single colony, whereas those for generation 8-10 were seeded using cultures from a different colony. 1,000 synchronized N2 L1 animals, 400 μl of 10x *E. coli* OP50-1 and a very low dose of *N. parisii* spores were combined in a centrifuge tube, evenly mixed, and pipetted onto an unseeded 6-cm NGM plate. Plates were incubated at 20°C for 72 hours. 10 gravid adult animals were then picked onto seeded 10-cm NGM plates with the desired bacteria. Every 4 days (representing a single generation), 10 gravid adult animals were passaged onto new seeded plates of the same bacteria while the rest of the plate was washed and fixed for imaging. At the 10^th^ passage, animals were grown for 4 days and then fixed and stained for imaging. At every generation, the pathogen load in at least 20 animals was measured (% DY96), as well as the fraction of the population containing *N. parisii* spores (% infected).

### Statistical analysis

All data analysis was performed in GraphPad prism 10.0. One-way ANOVA with post-hoc (Tukey test) was used for all experiments. Statistical significance defined as p < 0.05 with the exception of Figure 6a and S9b where significance was defined as p < 0.1.

## Supplementary

**Figure S1.**
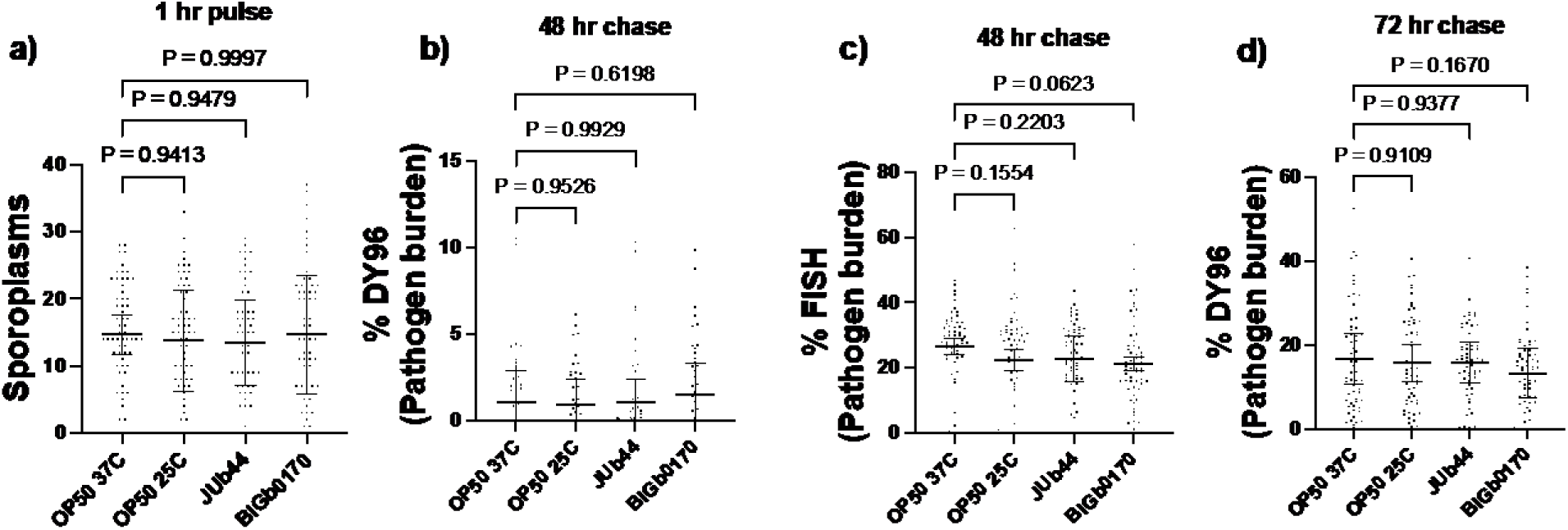
Growth on *C. scopthalmum* JUb44 and *S. multivorum* BIGb0170 for 24 hours does not impact *N. parisii* infection. (a-d) Synchronized N2 L1 animals were grown on bacterial lawns for 24 hours and then pulse infected with *N. parisii* and *E. coli* OP50 for one hour and fixed at 1 hpi (a), 48hpi (b,c), or 72hpi (d). Samples were stained with FISH probes and DY96 as indicated by the Y-axis in (b-d). Data is from three independent replicates of 20 worms each. Mean ± SD represented by horizontal bars. P-values determined via one-way ANOVA with post hoc.

**Figure S2.**
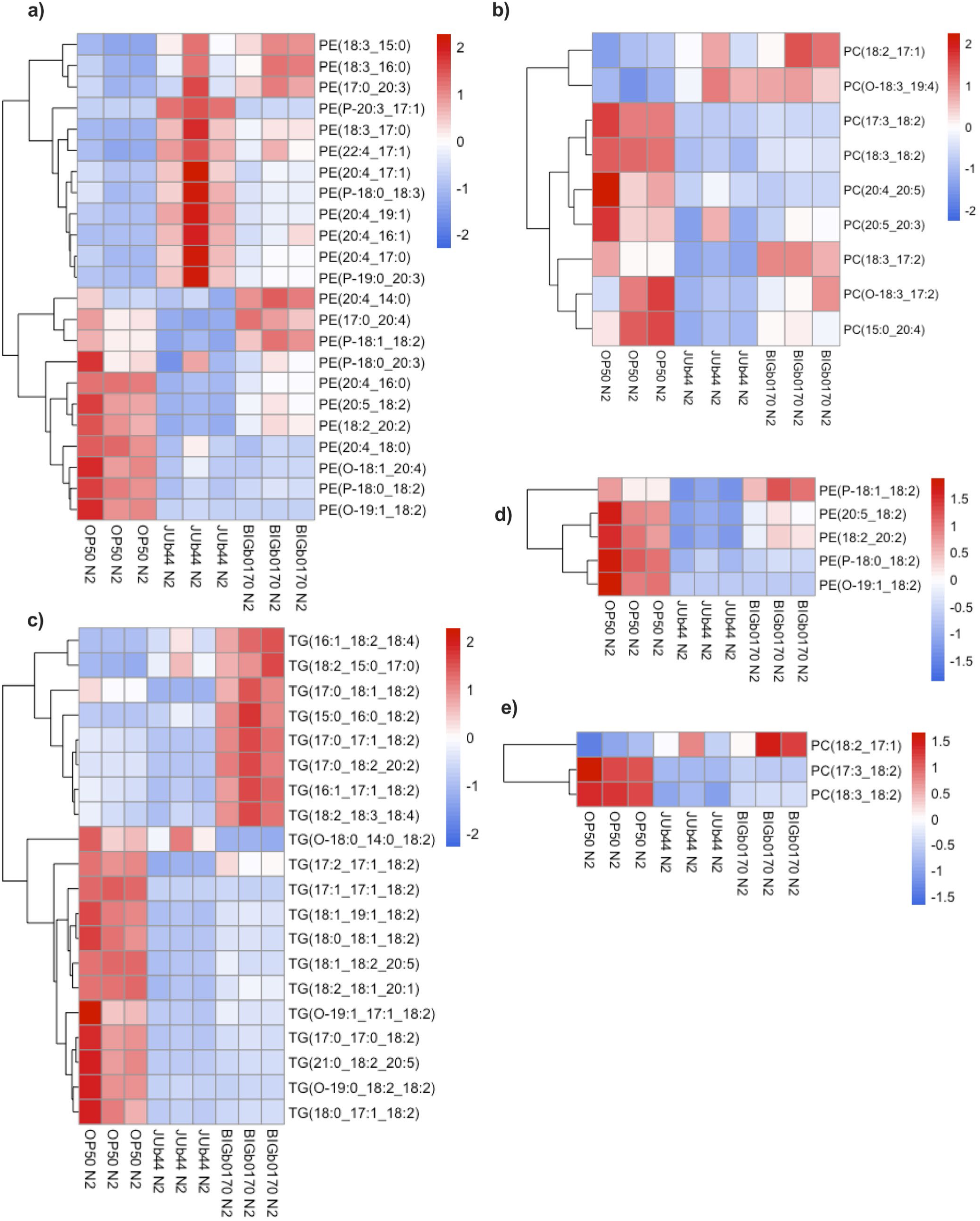
Abundance of triglycerides (TG), phosphatidylcholines (PC), and phosphatidylethanolamines (PE) in N2 animals fed *E. coli* OP50*, C. scopthalmum* JUb44, or *S. multivorum* BIGb0170. (a-b) Heat maps depicting the overall abundance of phosphatidylethanolamines (a) and phosphatidylcholines that contain at least one polyunsaturated fatty acid chain (b). (c-e) Heat maps depicting the abundance of 18:2 acyl chain containing triglycerides (c), phosphatidylethanolamines (d), and phosphatidylcholines (e). Data is from three independent replicates.

**Figure S3.**
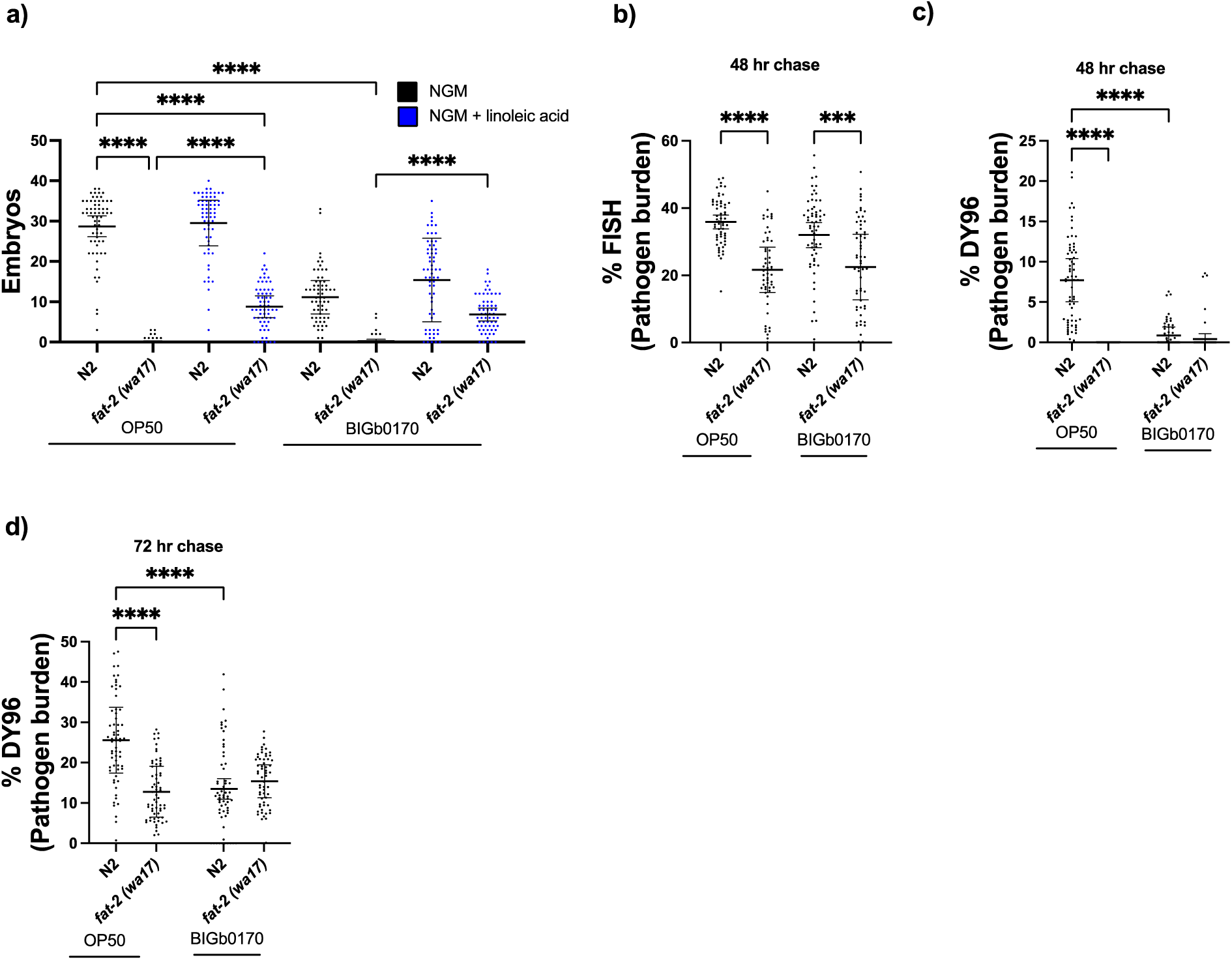
*fat-2 (wa17)* mutants display reduced *N. parisii* infection burdens. (a) N2 or *fat-2 (wa17)* animals were grown in the presence (blue) or absence (black) of linoleic acid for 72 hours and infected for one hour. The number of embryos was quantified. The bacterial diet is denoted with a solid line below the X-axis. (b-d) 72 hour old adult N2 or *fat-2 (wa17)* animals were infected for one hour with *N. parisii* and fixed either 48 (b,c) or 72 (d) hours post-infection. The level of meronts (b) and spores (c,d) were quantified. Data is from three independent replicates of 16-20 worms each. Mean ± SD represented by horizontal bars. P-values determined via one-way ANOVA with post hoc. Significance defined as *** p < 0.001, **** p < 0.0001.

**Figure S4.**
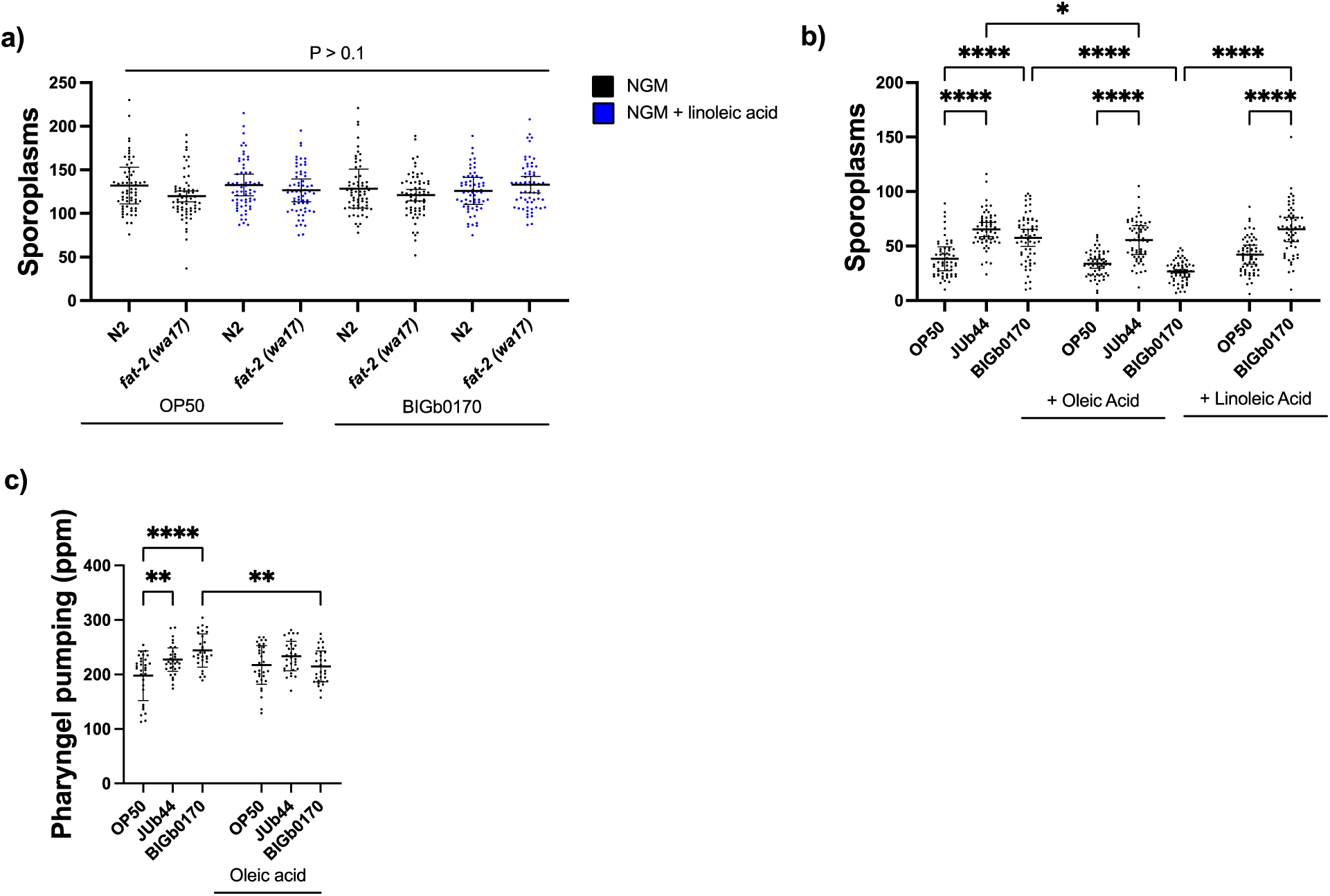
Oleic acid supplementation reduces pharyngeal pumping rates and *N. parisii* invasion on *S. multivorum* BIGb0170. (a) N2 or *fat-2 (wa17)* animals were grown in the presence (blue) or absence (black) of linoleic acid for 72 hours and infected for one hour. The number of sporoplasms was quantified. (b) N2 animals were grown on *E. coli* OP50, *C. scopthalmum* JUb44 or *S. multivorum* BIGb0170 in the presence or absence of oleic or linoleic acid for 72 hours (b,c) and infected for one hour (b). The number of sporoplasms (b) and pharyngeal pumping (c) was quantified. The supplemented fatty acids are denoted with a solid line below the X-axis. Data is from three independent replicates of 10 (c) or 20 worms each (a-b). Mean ± SD represented by horizontal bars. P-values determined via one-way ANOVA with post hoc. Significance defined as * p < 0.05, ** p < 0.01, **** p < 0.0001.

**Figure S5.**
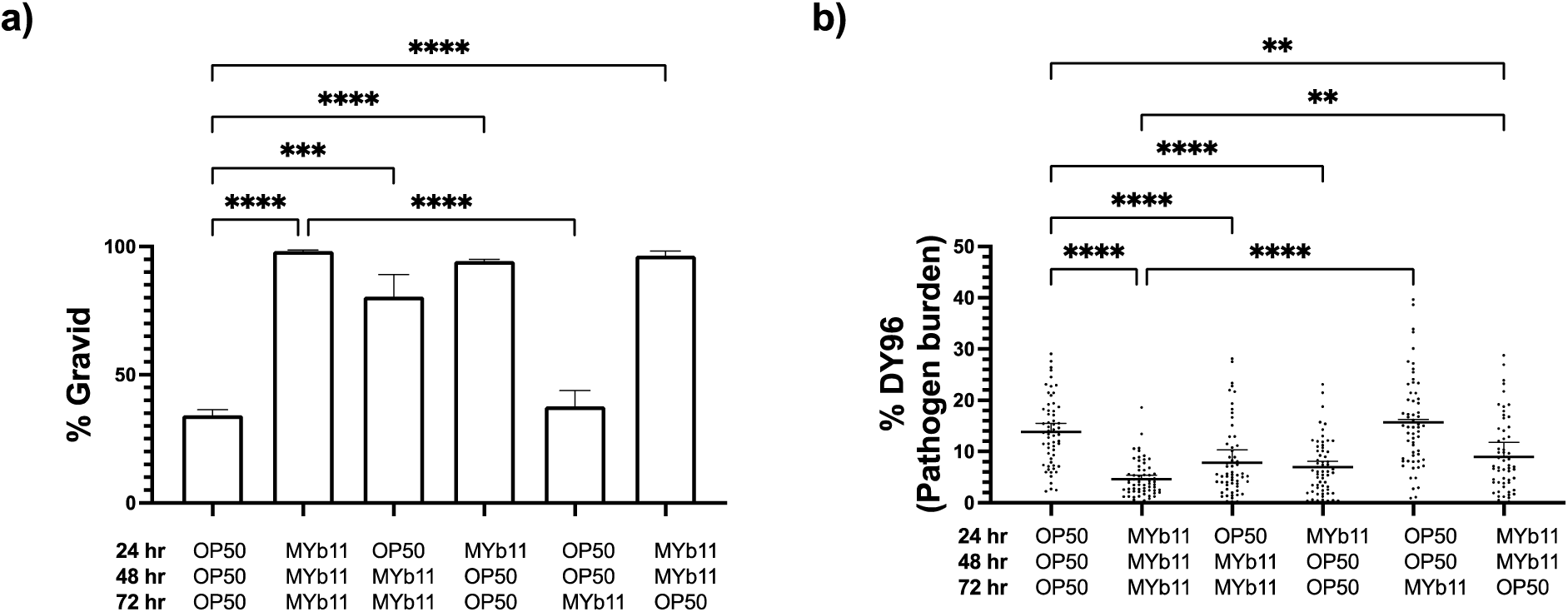
Protection by MYb11 against *N*. *parisii* can occur during the first 48 hours of exposure. (a-b) Animals were placed on diets of either *E. coli* OP50 or *P. lurida* MYb11 for varying amounts of time as indicated by the X-axis. Population fitness (a) and pathogen load (b) were quantified Data is from three independent replicates of at least 20 worms each (a-b) Mean ± SD represented by horizontal bars. P-values determined via one-way ANOVA with post hoc. Significance defined as ** p < 0.01, *** p < 0.001, **** p < 0.0001.

**Figure S6.**
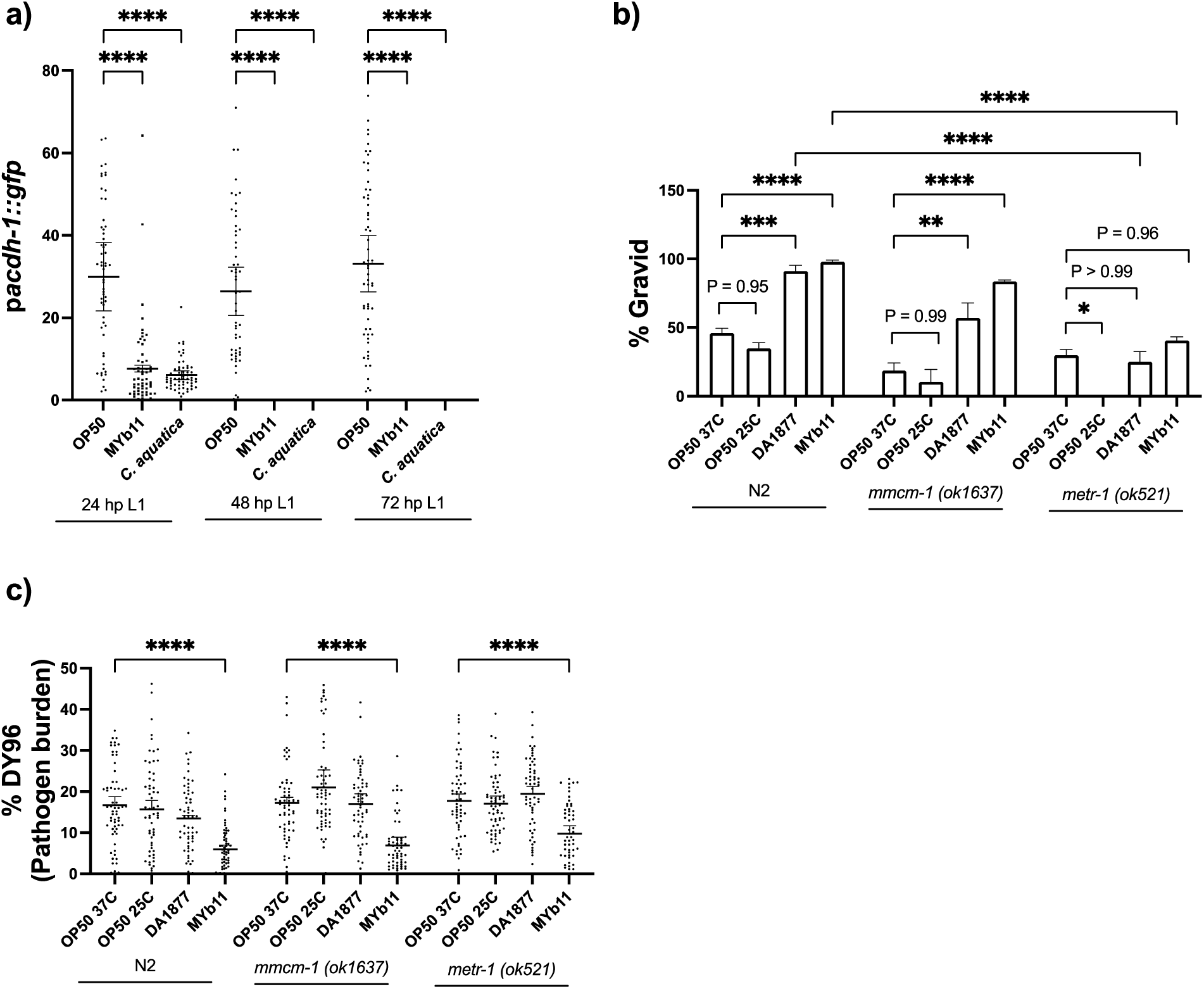
*P. lurida* MYb11 provides resistance against *N. parisii* which is independent of vitamin B12 developmental acceleration. (a) *acdh-1* expression levels were measured on various diets to assess the presence (GFP off) or absence (GFP on) of vitamin B12 in synchronized N2 animals every 24 hours. (b-c) Synchronized N2, *mmcm-1 (ok1637)* or *metr-1 (ok521)* L1’s were pulse infected with *N. parisii* for 1 hour on *E. coli* OP50 prior to washing and splitting the populations onto individual seeded plates. 72 hours later, population fitness (b) and pathogen load (c) were quantified. Data is from three independent replicates of at least 20 (a,c) or 100 worms each (b). Mean ± SD represented by horizontal bars. P-values determined via one-way ANOVA with post hoc. Significance defined as: ** p < 0.01, **** p < 0.0001.

**Figure S7.**
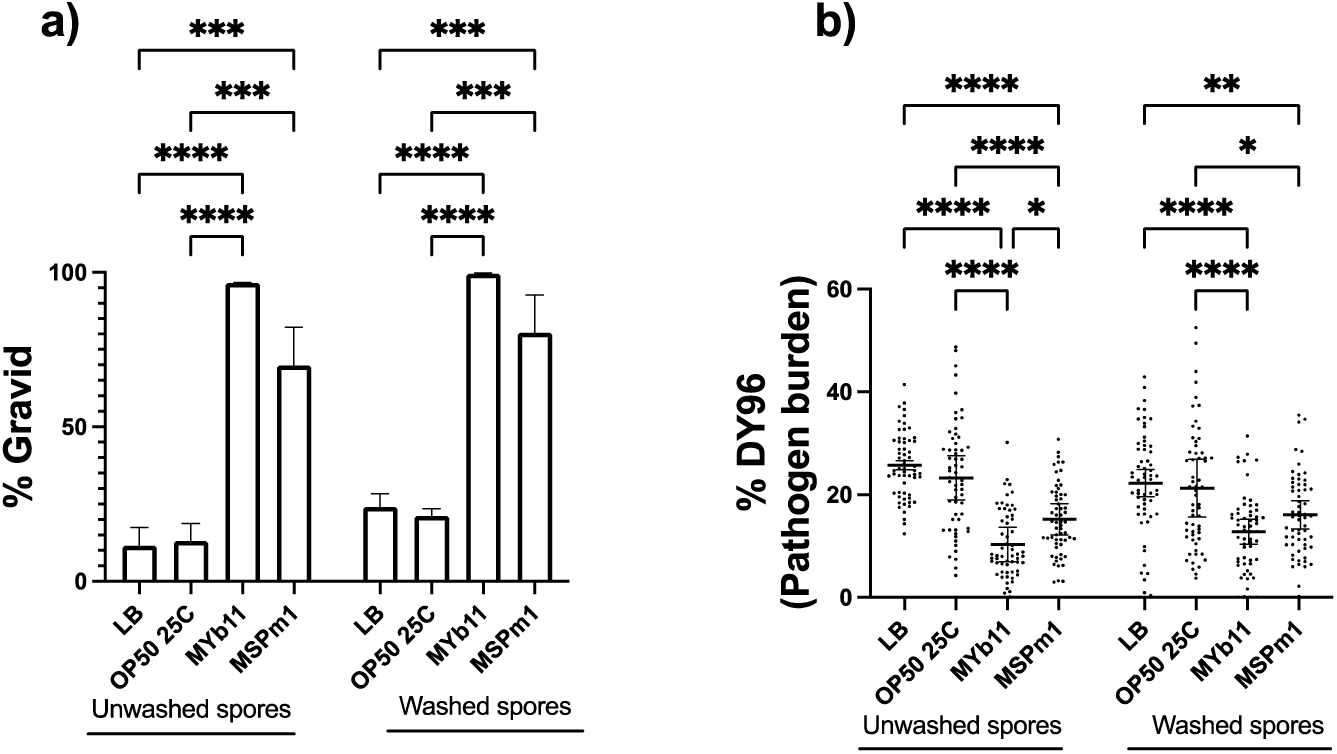
*P. lurida* MYb11 and *P. mendocina* MSPm1 secrete molecules which directly act on *N. parisii* spores. (a-b) Spores incubated in supernatants denoted on the X-axis were washed to remove residual supernatants prior to L1 pulse infection. Population fitness (a) or pathogen load (b) was quantified. Data is from three independent replicates of 100 worms (a) or 20 worms each (c). Mean ± SD represented by horizontal bars. P-values determined via one-way ANOVA with post hoc. Significance defined as * p < 0.05, ** p < 0.01, *** p < 0.001, **** p < 0.0001.

**Figure S8.**
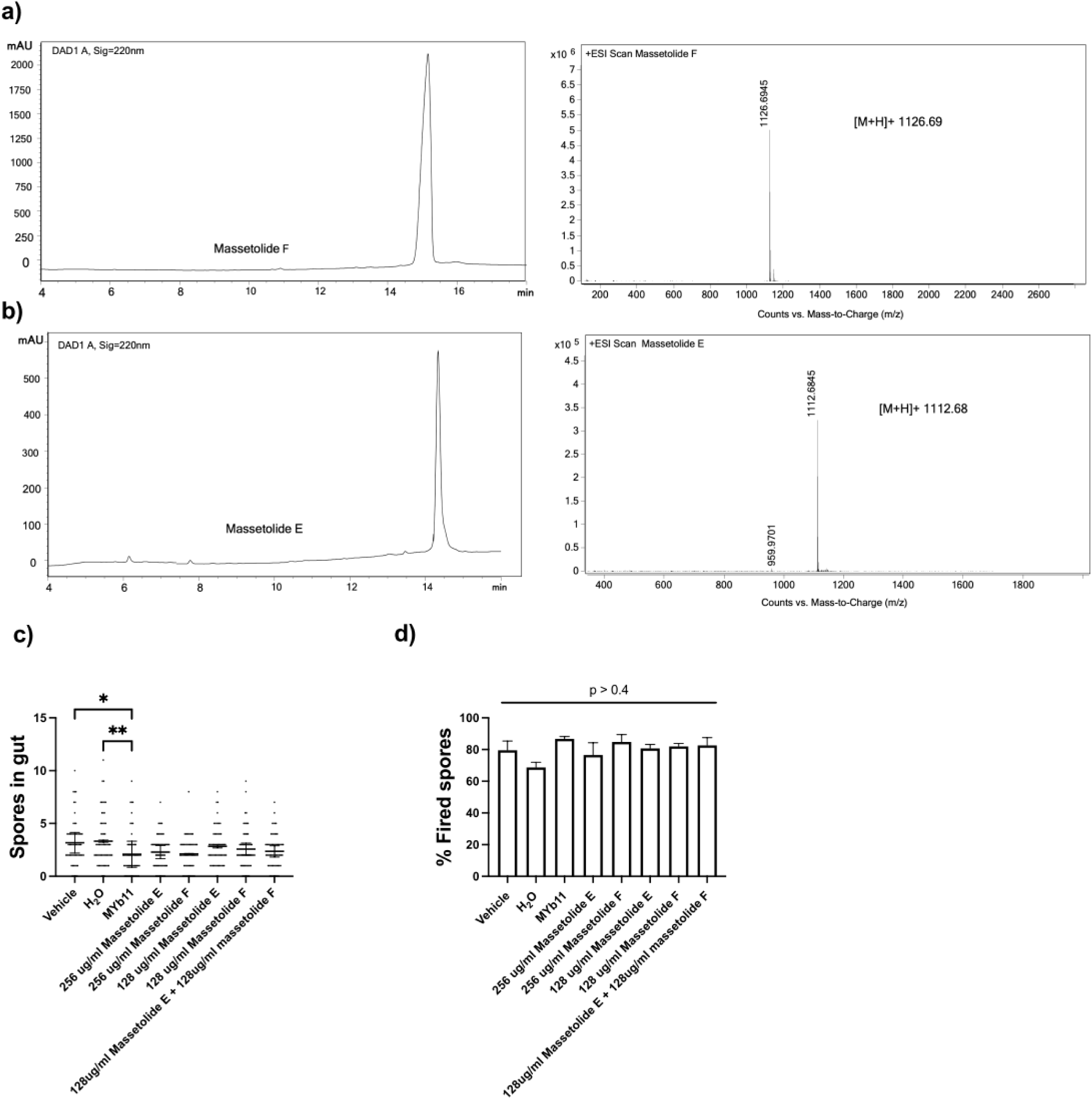
Massetolides E and F and their impact on the number of spores and the fraction of fired spores after in vitro incubation. (a-b) The HPLC chromatogram of massetolide F (a) and E (b) isolated from *P. lurida* MYb11 along with their high-resolution mass spectrometry (HR-MS) in positive ion mode. Massetolide F was identified by HR-MS as indicated by the peak at m/z 1126.69 [M+H]+ and massetolide E was identified by HR-MS as indicated by the peak at m/z 1112.68 [M+H]+. (c-d) Spores were incubated in either a vehicle control (0.5% DMSO), water, *P. lurida* MYb11 supernatant or massetolide E and/or F in various concentrations prior to L1 infection and the number of spores (c) and fraction of fired spores (d) were determined. Data is from three independent replicates of 20 worms each and at least 50 spores each (a-b). Mean ± SD represented by horizontal bars. P-values determined via one-way ANOVA with post hoc. Significance defined as * p < 0.05, ** p < 0.01.

**Figure S9.**
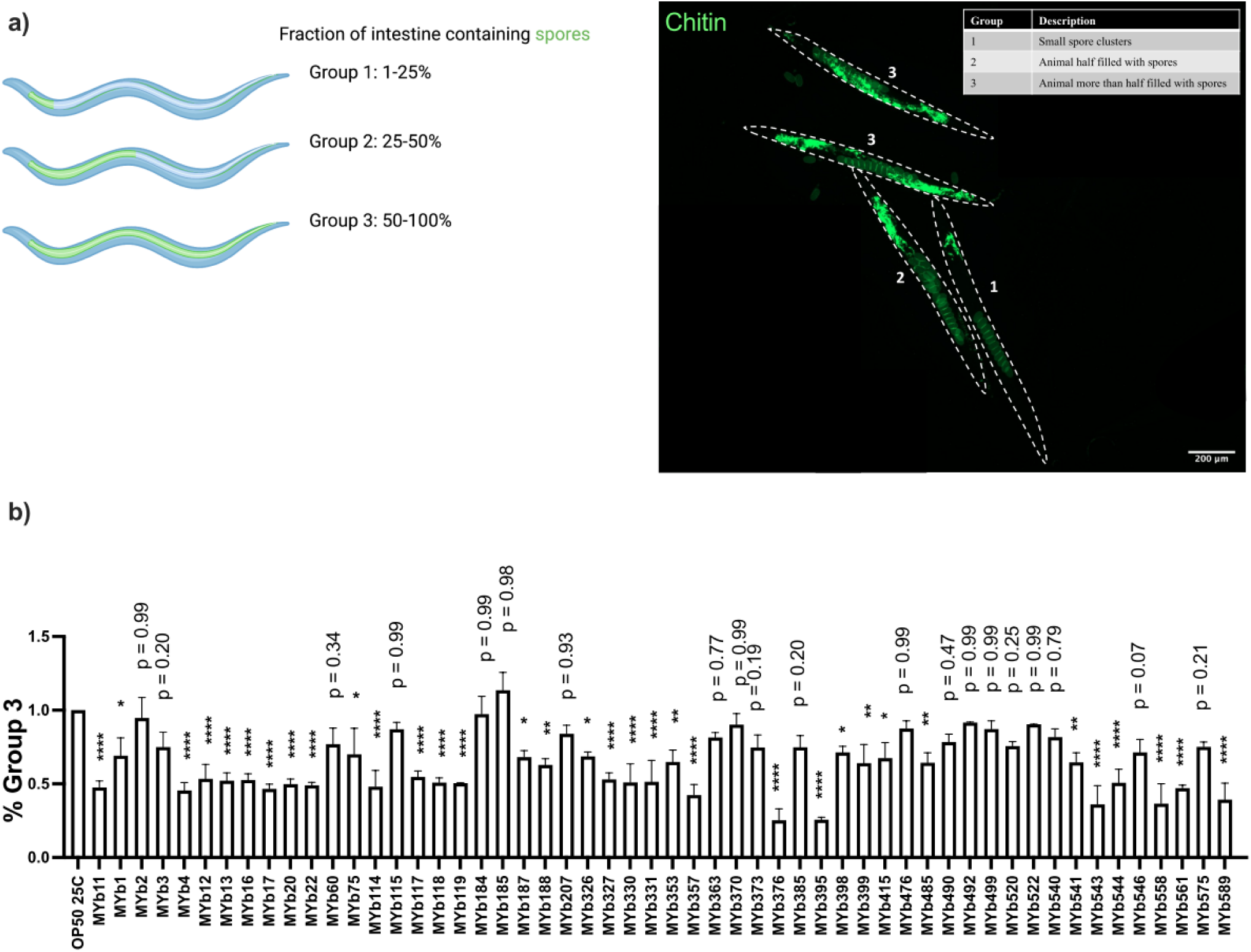
Cultured media from *Pseudomonas* spp. reduce *N. parisii* spore infectivity. (a) A schematic depicting the infection scale used to quantify pathogen burden (left) and a representative image of 72-hour old worms (outlined in dashed white lines) infected with *N. parisii* incubated in *P. lurida* MYb11 supernatant (right). Spores and embryos are stained with the chitin binding dye DY96. The number next to each nematode indicates the group under which it would be categorized using this infection scale. Scale bar represents 200 μm. Schematic generated via Biorender.com. (b) The percentage of animals belonging to group three is normalized relative to animals infected with spores incubated in *E. coli* OP50 supernatant. The X-axis denotes the supernatants in which *N. parisii* spores were incubated. Data is from three independent replicates of at least 50 worms each. Mean ± SD represented by horizontal bars. P-values determined via one-way ANOVA with post hoc. Significance defined as * p < 0.1, ** p < 0.01, **** p < 0.0001 and is relative to OP50 25°C.

**Figure S10.**
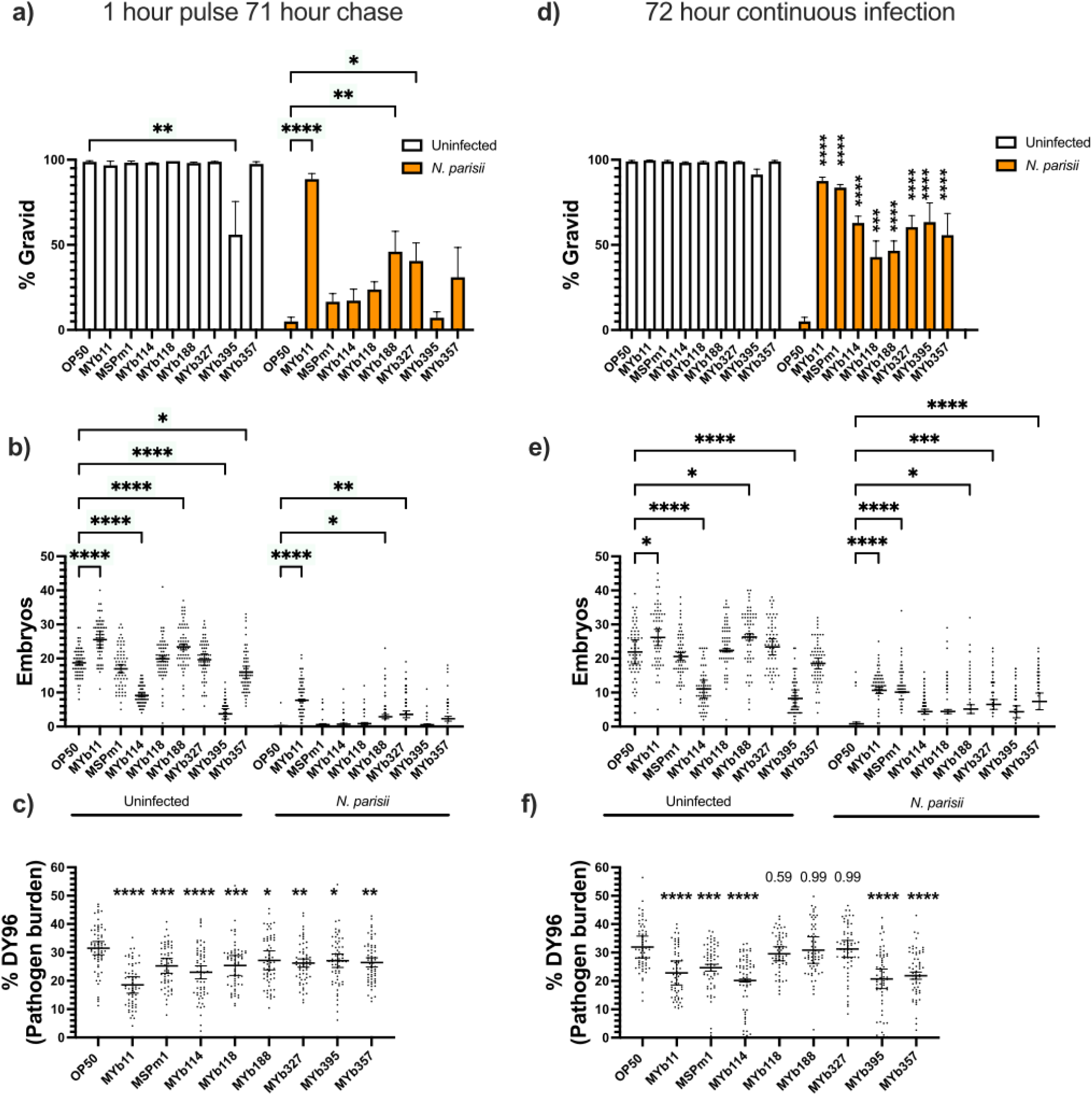
Infected animals grown on lawns of *Pseudomonas* spp. display improved fitness. Synchronized N2 L1 animals were pulse infected for one hour (a-c) or continuously infected for 72 hours (d-f). Population fitness (a,d), the number of embryos per worm (b,e) and the pathogen burden (c,f) are displayed. Data is from three independent replicates of at least 20 (b-c,e-f) or 100 (a,d) worms each. Mean ± SD represented by horizontal bars. P-values determined via one-way ANOVA with post hoc. Significance defined as * p < 0.05, ** p < 0.01, *** p < 0.001, **** p < 0.0001 and is relative to OP50 25°C of the corresponding condition.

**Figure S11.**
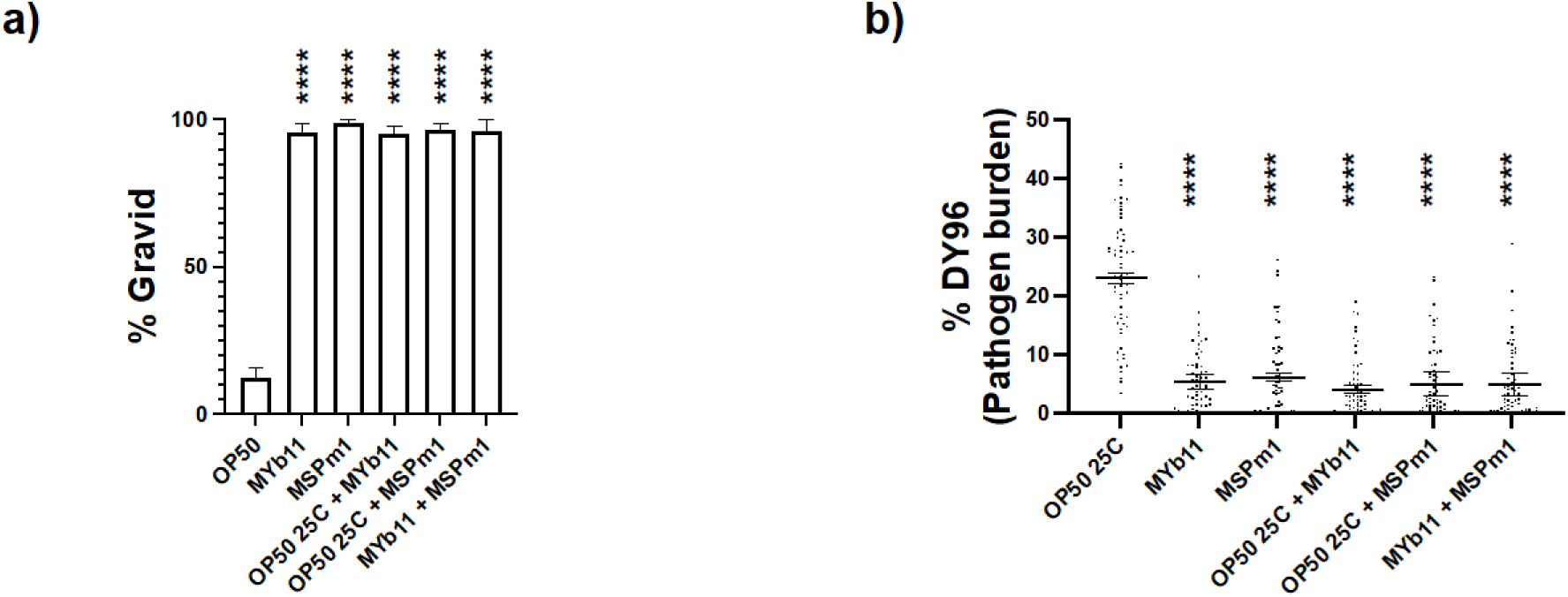
Infected animals grown on a combination of *P. lurida* MYb11 and *P. mendocina* MSPm1 result in decreased pathogen burden and improved host fitness. (a,b) Synchronized N2 L1 animals were continuously infected on lawns of bacteria indicated on the X-axis for 72 hours. Two bacterial species indicate a combination of the two in a 1:1 ratio (90 µl each) used to seed the NGM plates. The fraction of gravid adults (a) and the pathogen burden (b) are displayed. Data is from three independent replicates of at least 100 (a) or 20 (b) worms each. Mean ± SD represented by horizontal bars. P-values determined via one-way ANOVA with post hoc. Significance defined as **** p < 0.0001 and is relative to OP50 25°C of the corresponding condition.

**Table S1.**
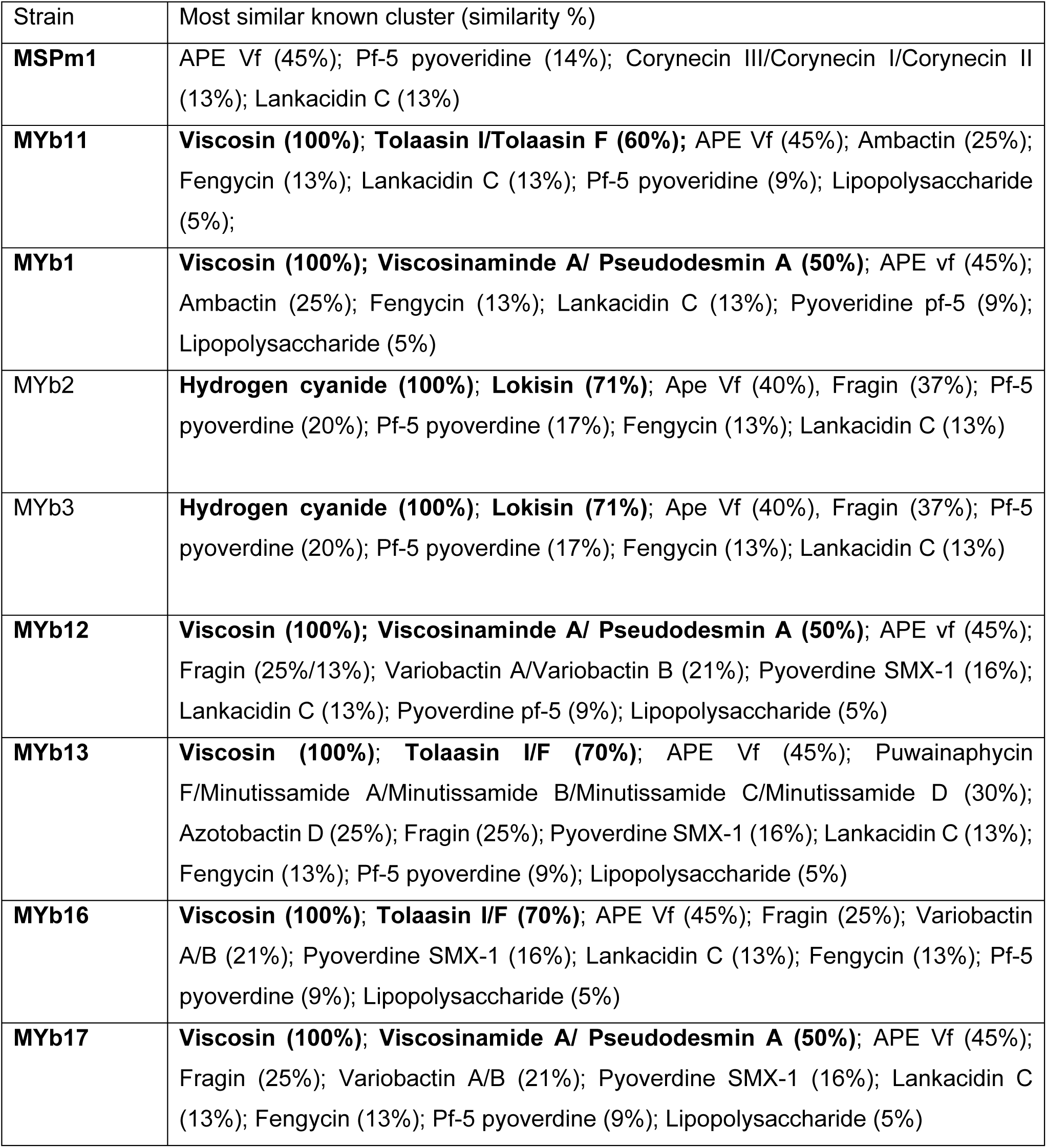
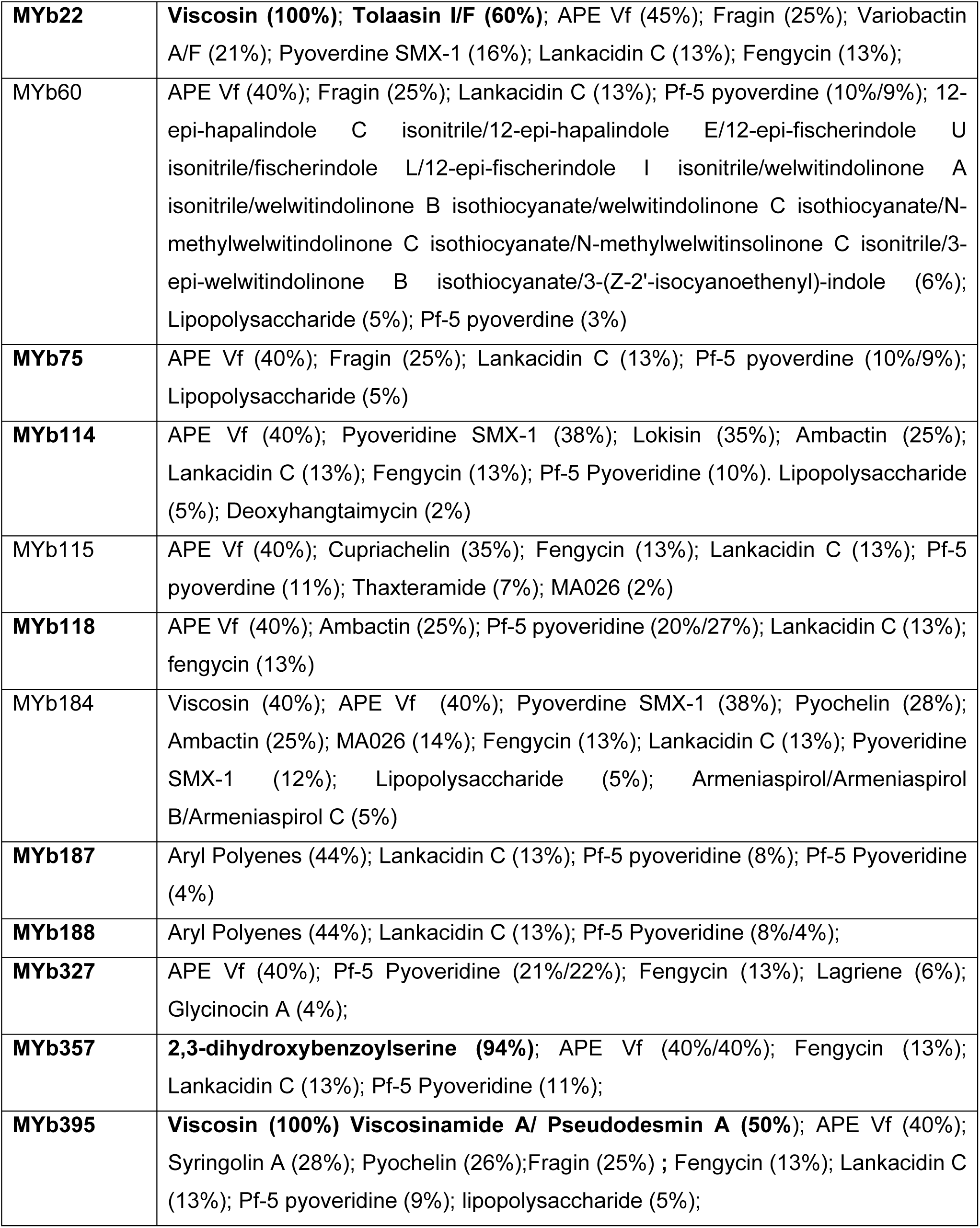
Diversity of natural compounds produced by *Pseudomonas* spp. The results from antiSMASH analysis are indicated under “Most similar known cluster”. A percentage in between brackets indicates the similarity score to the clusters. The indication of two different percentage scores for the same predicted cluster is representative of two separate clusters at different genomic regions. Similarities of 50% or higher are bolded. Strains with tested inhibitory activity against *N. parisii* are bolded.

| Strain | Most similar known cluster (similarity %) |
| --- | --- |
| **MSPm1** | APE Vf (45%); Pf-5 pyoverdine (14%); Corynecin III/Corynecin I/Corynecin II (13%); Lankacidin C (13%) |
| **MYb11** | **Viscosin (100%); Tolaasin I/Tolaasin F (60%);** APE Vf (45%); Ambactin (25%); Fengycin (13%); Lankacidin C (13%); Pf-5 pyoverdine (9%); Lipopolysaccharide (5%); |
| **MYb1** | **Viscosin (100%); Viscosinamide A/ Pseudodesmin A (50%);** APE vf (45%); Ambactin (25%); Fengycin (13%); Lankacidin C (13%); Pyoverdine pf-5 (9%); Lipopolysaccharide (5%) |
| MYb2 | **Hydrogen cyanide (100%); Lokisin (71%);** Ape Vf (40%), Fragin (37%); Pf-5 pyoverdine (20%); Pf-5 pyoverdine (17%); Fengycin (13%); Lankacidin C (13%) |
| MYb3 | **Hydrogen cyanide (100%); Lokisin (71%);** Ape Vf (40%), Fragin (37%); Pf-5 pyoverdine (20%); Pf-5 pyoverdine (17%); Fengycin (13%); Lankacidin C (13%) |
| **MYb12** | **Viscosin (100%); Viscosinamide A/ Pseudodesmin A (50%);** APE vf (45%); Fragin (25%/13%); Variobactin A/Variobactin B (21%); Pyoverdine SMX-1 (16%); Lankacidin C (13%); Pyoverdine pf-5 (9%); Lipopolysaccharide (5%) |
| **MYb13** | **Viscosin (100%); Tolaasin I/F (70%);** APE Vf (45%); Puwainaphycin F/Minutissamide A/Minutissamide B/Minutissamide C/Minutissamide D (30%); Azotobactin D (25%); Fragin (25%); Pyoverdine SMX-1 (16%); Lankacidin C (13%); Fengycin (13%); Pf-5 pyoverdine (9%); Lipopolysaccharide (5%) |
| **MYb16** | **Viscosin (100%); Tolaasin I/F (70%);** APE Vf (45%); Fragin (25%); Variobactin A/B (21%); Pyoverdine SMX-1 (16%); Lankacidin C (13%); Fengycin (13%); Pf-5 pyoverdine (9%); Lipopolysaccharide (5%) |
| **MYb17** | **Viscosin (100%); Viscosinamide A/ Pseudodesmin A (50%);** APE Vf (45%); Fragin (25%); Variobactin A/B (21%); Pyoverdine SMX-1 (16%); Lankacidin C (13%); Fengycin (13%); Pf-5 pyoverdine (9%); Lipopolysaccharide (5%) |
| **MYb22** | **Viscosin (100%); Tolaasin I/F (60%);** APE Vf (45%); Fragin (25%); Variobactin A/F (21%); Pyoverdine SMX-1 (16%); Lankacidin C (13%); Fengycin (13%); |
| MYb60 | APE Vf (40%); Fragin (25%); Lankacidin C (13%); Pf-5 pyoverdine (10%/9%); 12-epi-hapalindole C isonitrile/12-epi-hapalindole E/12-epi-fischerindole U isonitrile/fischerindole L/12-epi-fischerindole I isonitrile/welwitindolinone A isonitrile/welwitindolinone B isothiocyanate/welwitindolinone C isothiocyanate/N-methylwelwitindolinone C isothiocyanate/N-methylwelwitinsolinone C isonitrile/3-epi-welwitindolinone B isothiocyanate/3-(Z-2'-isocyanoethenyl)-indole (6%); Lipopolysaccharide (5%); Pf-5 pyoverdine (3%) |
| **MYb75** | APE Vf (40%); Fragin (25%); Lankacidin C (13%); Pf-5 pyoverdine (10%/9%); Lipopolysaccharide (5%) |
| **MYb114** | APE Vf (40%); Pyoveridine SMX-1 (38%); Lokisin (35%); Ambactin (25%); Lankacidin C (13%); Fengycin (13%); Pf-5 Pyoveridine (10%). Lipopolysaccharide (5%); Deoxyhangtaimycin (2%) |
| MYb115 | APE Vf (40%); Cupriachelin (35%); Fengycin (13%); Lankacidin C (13%); Pf-5 pyoverdine (11%); Thaxteramide (7%); MA026 (2%) |
| **MYb118** | APE Vf (40%); Ambactin (25%); Pf-5 pyoveridine (20%/27%); Lankacidin C (13%); fengycin (13%) |
| MYb184 | Viscosin (40%); APE Vf (40%); Pyoverdine SMX-1 (38%); Pyochelin (28%); Ambactin (25%); MA026 (14%); Fengycin (13%); Lankacidin C (13%); Pyoveridine SMX-1 (12%); Lipopolysaccharide (5%); Armeniaspirol/Armeniaspirol B/Armeniaspirol C (5%) |
| **MYb187** | Aryl Polyenes (44%); Lankacidin C (13%); Pf-5 pyoveridine (8%); Pf-5 Pyoveridine (4%) |
| **MYb188** | Aryl Polyenes (44%); Lankacidin C (13%); Pf-5 Pyoveridine (8%/4%); |
| **MYb327** | APE Vf (40%); Pf-5 Pyoveridine (21%/22%); Fengycin (13%); Lagriene (6%); Glycinocin A (4%); |
| **MYb357** | **2,3-dihydroxybenzoylserine (94%);** APE Vf (40%/40%); Fengycin (13%); Lankacidin C (13%); Pf-5 Pyoveridine (11%); |
| **MYb395** | **Viscosin (100%) Viscosinamide A/ Pseudodesmin A (50%);** APE Vf (40%); Syringolin A (28%); Pyochelin (26%);Fragin (25%) ; Fengycin (13%); Lankacidin C (13%); Pf-5 pyoveridine (9%); lipopolysaccharide (5%); |

**Table S2.**
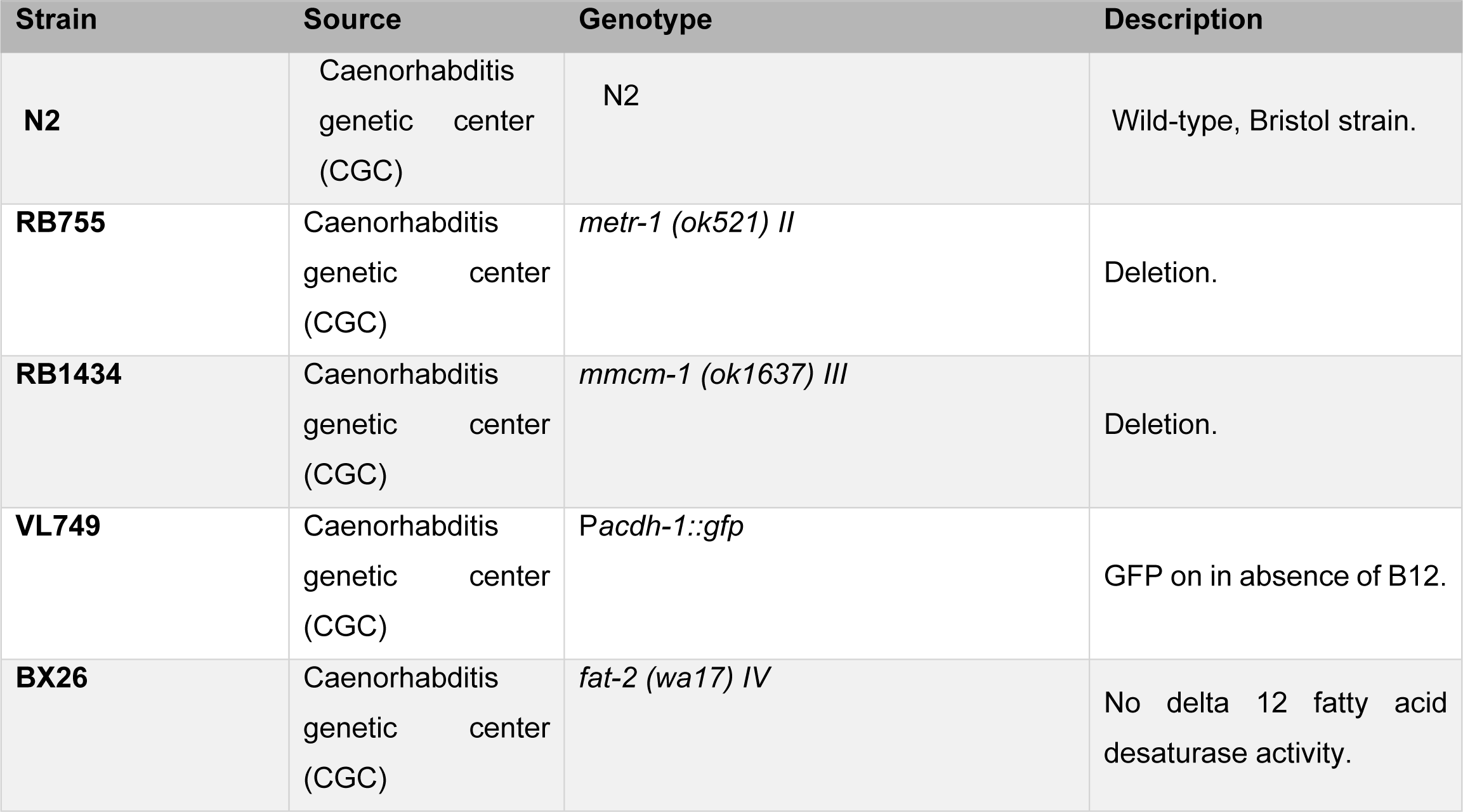
List of *C. elegans* strains used in this study.

**Table S3.**
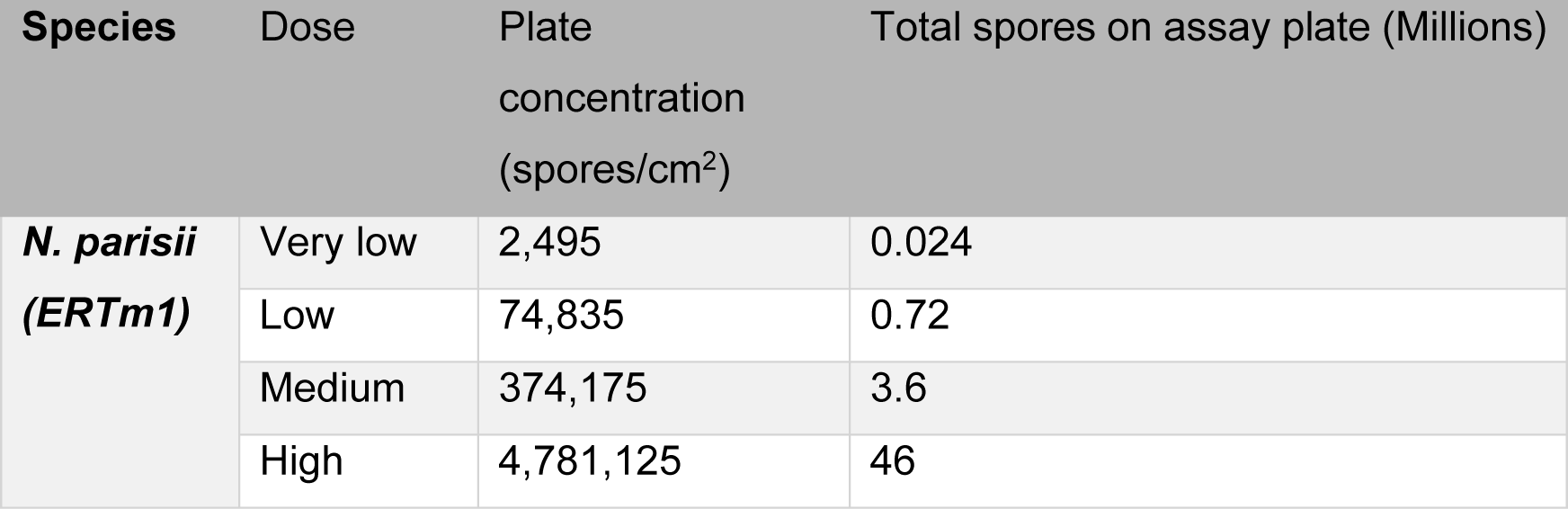
Spore doses used in this study. The varying doses of spores (as defined in methods) are listed. Plate concentration refers to the number of spores occupied per cm^2^ on a 6-cm NGM plate. The total number of spores present on a single assay plate are listed for the various doses in millions of spores.

